# Characterisation of Two *Plasmodium* Virulence Factors Important for Lipid Metabolism and Disease Progression *In Vivo*

**DOI:** 10.64898/2025.12.21.695507

**Authors:** Martina S. Paoletta, Thorey K. Jonsdottir, Alison Kemp, Sophia Hernandez, Scott A. Chisholm, María Rayón Díaz, Alexandra Berg, Vikash Pandey, Cyrille Y. Botté, Ronnie P-A. Berntsson, Ross F. Waller, Julian C. Rayner, Ellen S.C. Bushell

## Abstract

Malaria is caused by *Plasmodium* parasites and claims 610,000 deaths annually. Parasite invasion of red blood cells (RBCs) involves secretion of multiple proteins from specialised apical organelles (micronemes, rhoptries, and dense granules). These collectively facilitate host cell entry, establish a protective parasitophorous vacuole (PV), and directly and indirectly mediate interactions between the infected RBC (iRBC) and the endothelial wall. The cytoadherence of iRBCs results in sequestration within organs and plays a critical role in pathogenesis and virulence, but the function of many secreted parasite proteins remains poorly characterised. Here, we leverage spatial proteomics from *Plasmodium falciparum* to identify two novel *Plasmodium berghei* orthologue proteins containing hydrolase domains (MAP1, PBANKA_1425900, and RhoSH, PBANKA_1001500). Ultrastructure expansion microscopy reveals their localisation to the rhoptries in late schizogony, where co-immunoprecipitation assays show they interact with each other. *In vivo* studies demonstrate that these proteins help the parasite evade spleen-mediated clearance and contribute to disease progression. Deletion of both genes disrupts lipid metabolism and PV membrane morphology, suggesting a role in establishing or maintaining the protective PV membrane. Loss of *map1* reduces sequestration of *P. berghei* iRBCs to adipose tissue, and conditional knockdown of the *P. falciparum map1* orthologue results in reduced CD36-mediated cytoadhesion, suggesting a cross-species role in sequestration and cytoadherence. Our findings identify MAP1 and RhoSH as key mediators of *Plasmodium* virulence.

## Introduction

Malaria is a life-threatening disease caused by *Plasmodium* parasites, which are transmitted through mosquito bites, resulting in 282 million cases and 610,000 deaths annually (World Health Organization, 2025). Morbidity and mortality in malaria arise from the parasite’s multiplication within the red blood cells (RBCs) of the vertebrate host.

The invasion of RBCs by *Plasmodium* involves the release of proteins from specialised apical secretory organelles that facilitate the initial attachment of the parasite to the host cell and contribute to the formation of the parasitophorous vacuole (PV). Proteins secreted from micronemes and the rhoptry neck support motility and early steps of host cell engagement, facilitating parasite entry. Proteins and lipids released from the rhoptry bulb and dense granules help to establish the PV membrane (PVM) and mediate early host-parasite interactions to support intracellular replication (Goldberg & Zimmerberg, 2020; Kats et al., 2006; Liffner et al., 2021).

Once infection is established, the parasite evades immune detection by promoting sequestration of the infected RBC (iRBC) to the microvasculature and thereby avoiding splenic clearance (Dondorp et al., 2000; Goldberg & Cowman, 2010; Maier et al., 2008; Spillman et al., 2015; Wahlgren et al., 2017). This is in part achieved by altering the iRBC cytoskeleton to increase rigidity and enhancing cytoadherence mediated by CD36 receptors in the endothelial cells (Rowe et al., 2009; Sherman et al., 2003). Parasite development inside the iRBC also necessitates extensive membrane biosynthesis and lipid metabolism to support parasite proliferation, which requires nutrient uptake, organelle replication, and expansion of the PVM and parasite plasma membrane (PPM) (Shunmugam et al., 2022; Vial et al., 2003). This is reflected in an increase of phospholipid (PL) content by ~600% upon *P. falciparum* RBC infection, upregulation of parasite lipid synthetic enzymes and trafficking of lipid manipulating enzymes such as phospholipases to the iRBC membranes and host-parasite interface (Liu et al. 2024, Sheokand et al., 2023 Vial et al., 2003). The fatty acids and polar heads for parasite membrane lipids are obtained by a combination of *de novo* synthesis and scavenging, where reassembly of components from different sources is increasingly recognised in its importance for supporting parasite intraerythrocytic development (Shunmugam et al., 2022).

The rodent malaria parasite *Plasmodium berghei* serves as a genetically tractable *in vivo* model that has been experimentally validated for studying virulence factors and their roles in malaria pathology. The translocon machinery and protein sorting hubs (e.g., intra-erythrocytic *P. berghei*-induced structures and Maurer’s clefts) underlying virulence are largely conserved across rodent and human malaria parasites (De Niz et al., 2016), making this model suitable for investigating protein functions in the context of organ-specific interactions and host immune responses.

Here, we characterised two conserved malaria parasite virulence factors using the *P. berghei in vivo* model: Merozoite Attachment Protein 1 (MAP1, PBANKA_1425900) and Rhoptry Serine Hydrolase (RhoSH, PBANKA_1001500). MAP1 and RhoSH co-precipitate each other according to immunoprecipitation data during late schizogony, where both proteins are localised to the rhoptries. Both MAP1 and RhoSH are indirectly involved in evading spleen-mediated clearance of iRBCs, with MAP1 in particular playing a role in sequestration to adipose tissue. Loss of either gene also disrupts lipid content and homeostasis, consistent with a role in establishing or maintaining the PV/PVM. Conditional knockdown of the PfMAP1 ortholog in the major human malaria pathogen *P. falciparum* has no effect on growth *in vitro,* but results in reduced cytoadhesion of iRBCs to the major endothelial receptor CD36, suggesting a conserved cross-species role in sequestration and virulence.

## Results

### 1. Identification and structural insights into novel RBC-remodelling proteins

Using a spatial proteomics dataset based on density gradient separation of subcellular compartments (Chisholm et al., 2025), we identified previously uncharacterised proteins of *Plasmodium falciparum* schizonts that cluster with exported proteins or those residing within the PV and PVM. To prioritise candidates, we evaluated each gene’s importance for blood-stage growth based on the *piggyBac* insertional mutagenesis screen in *P. falciparum* (Zhang et al., 2018), and identified orthologues in *P. berghei* to facilitate *in vivo* studies.

We selected the α/β hydrolase PBANKA_1001500 (RhoSH/S9C) for characterisation as its *P. falciparum* orthologue (PF3D7_1120400) was predicted to be exported, and its knockout in *P. berghei* causes slow growth during blood-stage infection in mice (Bushell et al., 2017). We also selected PBANKA_1425900 (MAP1), the syntenic orthologue of PF3D7_0811600, which was localised to the PV cluster. Both proteins were non-mutable in the *piggyBac* transposon screen during the blood stages in *P. falciparum,* with MAP1 found to be non-mutable in *P. knowlesi* as well, suggesting they are essential during the blood stages (Elsworth et al., 2025; Oberstaller et al., 2025; Zhang et al., 2018). The knockout of MAP1 in *P. berghei* also results in a slow growth blood stage phenotype (Bushell et al., 2017). MAP1 orthologues are exclusive to Apicomplexa (ortholog group OG7_0046315; OrthoMCL database, release 7.0 (Fischer et al., 2011)) but lack functional annotation. MAP1 was also recently characterised as a heparin-binding protein that localises to the merozoite surface and is hypothesised to be involved in merozoite attachment during the invasion process (Gao et al., 2023), necessitating validation of the PVM localisation predicted by spatial proteomics.

We employed AlphaFold2 for structural predictions of RhoSH and MAP1, complemented by Foldseek for structural comparisons. Consistent with previous studies (Davison et al., 2022; Elahi et al., 2019), the AlphaFold2 model of RhoSH contains a well-predicted α/β hydrolase domain at the N-terminal region **(Fig. 1a, Fig. S1a)**. The remaining two-thirds of the protein model exhibited low to very low confidence and appeared predominantly disordered. Superimposition of the predicted RhoSH α/β hydrolase domain with structurally related, experimentally solved proteins retrieved through Foldseek (E-value < 1e^-10^) identified a serine hydrolase catalytic triad formed by Ser125-His218-Asp189.

**Figure 1:**
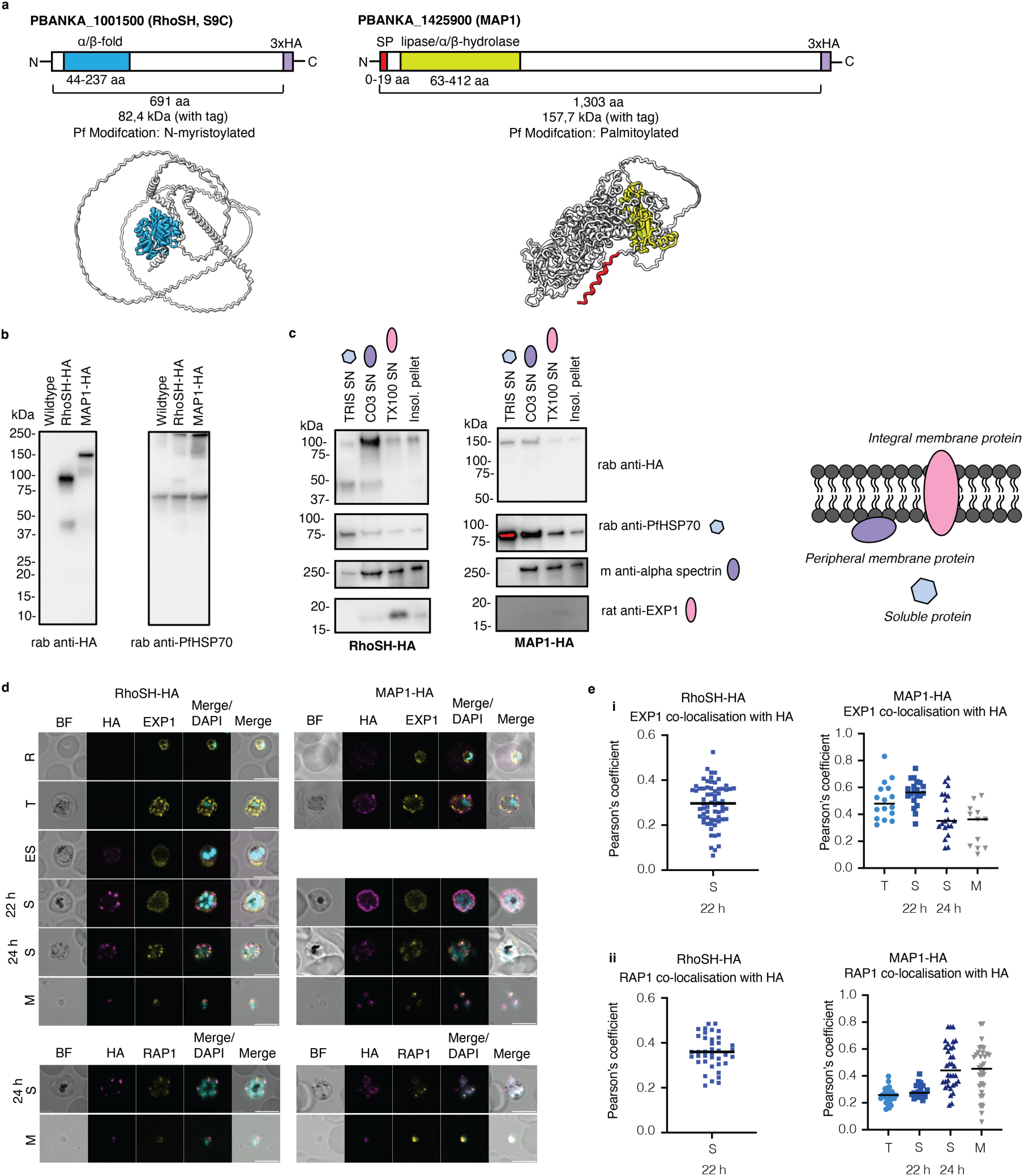
The expression and localisation of RhoSH and MAP1 during the intraerythrocytic cycle. **a** Schematic representation and the predicted AlphaFold2 models of RhoSH (left) and MAP1 (right). The identified domains are highlighted in blue (RhoSH) and yellow (MAP1) and the predicted signal peptide for MAP1 is marked in red in both the schematic and AlphaFold2 predicted structure. The schematics additionally include the C-terminal 3xHA epitope tag in purple. **b** Western blots prepared from schizont stage parasites purified using a Percoll density gradient, probed with mouse anti-HA (tagged proteins) and rabbit anti-PfHSP70 (loading control), where wildtype parasites serve as a negative control for the HA tag. **c** To assess the association of RhoSH and MAP1 with membranes, saponin-lysed schizont-enriched parasite pellets underwent serial sequential lysis in different buffers. Pellets were first lysed in hypotonic buffer (TRIS), then sodium carbonate and lastly Triton X-100. Supernatant (SN) was harvested after each treatment and the Triton X-100 insoluble pellet at the end. Rabbit anti-HA was used to probe for the target proteins, rabbit anti-PfHSP70 was used as a control for soluble proteins, mouse anti-α spectrin for peripherally membrane-associated proteins and rat anti-EXP1 for integral membrane proteins. Representative of three biological replicates. **d** An *ex vivo* culture was prepared from blood collected from heartbleed (ring stage) and samples collected for immunofluorescence assays at different time points over 24 hours. Rabbit anti-HA (magenta) was used to probe for the target protein, rat anti-EXP1 (yellow) used as a marker for the PVM and dense granules, RAP1 (yellow) as a marker for the rhoptries and DAPI (4′,6-diamidino-2-phenylindole) (cyan) to stain the nucleus. Scale bar = 5 µm. Representative of three biological replicates. **e** Pearson’s correlation coefficient was used to determine co-localisation of target proteins with either **i** EXP1 (PVM) or **ii** RAP1 (rhoptry bulb) at different stages from three independent experiments. T = trophozoite stage, ES = early schizont stage, S = schizont stage, either 22 h or 24 h, M = merozoite stage.

The AlphaFold2 prediction for MAP1 is, overall, of high confidence **(Fig. S1b)**. However, structural alignment searches using Foldseek did not yield any significant structural matches, which is consistent with the OrthoMCL database that suggests this may represent a structurally novel protein family within Apicomplexa. The only potential alignment was for the N-terminal region (∼residues 25-215), which showed structural similarity to a serine hydrolase domain, suggesting it could be a carboxylesterase or lipase. However, these matches had E-values ranging from 1.55e^-1^ to 6.39e^-1^, indicating low confidence in the prediction. The majority of lipases are α/β fold serine hydrolases, and although MAP1 appears to lack a canonical serine hydrolase catalytic triad, a potential non-canonical Cys194-Ser625-His193 triad is present (Arpigny & Jaeger, 1999; Farooqui et al., 1987). Additionally, a potential signal peptide (SP) has been predicted for MAP1, spanning residues 1-19 **(Fig. 1a)**.

### 2. RhoSH and MAP1 are peripherally associated with rhoptry membranes in mature schizonts

Both *P. berghei* proteins (RhoSH and MAP1) were tagged with 3xHA at the C-terminal end using *Plasmo*GEM vectors (Bushell et al., 2017; Schwach et al., 2015), which integrate the epitope tag into the endogenous copy of the gene **(Fig. 1a).** Successful integration was confirmed by genotyping PCRs **(Fig. S2a-b)**. We confirmed tagging and expression of the tagged proteins by Western blot **(Fig. 1b)**. RhoSH-3xHA ran slightly higher than expected, likely due to post-translational modification of the protein or its association with membranes (Schlott et al., 2021). We also observed that RhoSH is processed, where a ∼45 kDa product containing the C-terminal tag was detected in the Western blot. MAP1-3xHA was mostly observed as a full-length protein, with two faint products observed just above 100 kDa, suggesting MAP1 is cleaved as well **(Fig. 1b, S3a)**.

Next, we biochemically characterised the membrane association of both proteins at the schizont stage, using a sequential solubility assay. We found that the majority of RhoSH-3xHA protein was solubilised using carbonate extraction, and a similar pattern was observed for MAP1-3xHA **(Fig. 1c)**. RhoSH was partially soluble using hypotonic lysis, whereas MAP1 was more soluble under these conditions. For both proteins, some fraction was released during Triton X-100 treatment (integral membrane proteins) whilst some remained insoluble **(Fig. 1c)**. These assays support a model where RhoSH and MAP1 are peripherally associated with membranes. Given that neither protein contains predicted transmembrane domains, this interaction is likely mediated by protein-protein interactions or by post-translational lipidation, which are known to mediate membrane binding, and which have been shown experimentally for their *P. falciparum* orthologues (palmitoylation in PfMAP1 and N-myristoylation in PfRhoSH) (Jones et al., 2012; Schlott et al., 2021; Wright et al., 2014).

We then investigated protein localisation and expression during the intraerythrocytic cell cycle by indirect immunofluorescence assays (IFAs). RhoSH-3xHA was predominantly observed in the schizont stages and free merozoites **(Fig. 1d)**. In schizonts, RhoSH showed an apical, punctuate pattern suggestive of localisation to apical secretory organelles, though it only partially co-localised with the rhoptry bulb protein RAP1 (Howard & Reese, 1990). In a subset of early schizonts, the protein was observed in distinct puncta partially overlapping with the PVM marker EXP1 (Hall et al., 1983), which might represent early rhoptry precursors or trafficking intermediates during the *de novo* formation of rhoptries in schizogony **(Fig. 1d-e)**. MAP1-3xHA was detectable throughout the asexual blood-stage cycle. A faint signal was observed within the RBC during the ring stage, while in trophozoites the protein was predominantly localised to the PVM, partially overlapping with EXP1. In mature schizonts and free merozoites, MAP1 accumulated at the apical end of the parasite where it co-localised with the rhoptry bulb marker RAP1 **(Fig. 1d-e)**. No HA signal was detected in the wild type (WT) control, as expected **(Fig. S3b)**.

Ultrastructure expansion microscopy (U-ExM) was used to better resolve the apical localisation of RhoSH and MAP1 in late schizonts **(Fig. 2)**. Both proteins overlapped with lipid-rich (BODIPY TRc) and protein-rich (N-hydroxysuccinimide (NHS) ester) regions of the rhoptries, previously described by Liffner et al., 2023 **(Fig. 2a-b)**. MAP1 was observed at the PVM in younger schizonts, in agreement with IFA results **(Fig. 2a).** MAP1 was occasionally detected at the periphery of merozoites where, contrary to previous reports (Gao et al., 2023), it appeared to be on the inner side of the BODIPY membrane stain rather than on the surface. MAP1 showed clear co-localisation with the rhoptry bulb marker RAP1, whereas RhoSH displayed more of a punctate distribution around the bulb and often concentrated at the neck, consistent with a rhoptry membrane localisation **(Fig. 2c)**. Neither MAP1 nor RhoSH co-localised with EXP1 in the dense granules during late schizont stages **(Fig. 2d)**.

**Figure 2:**
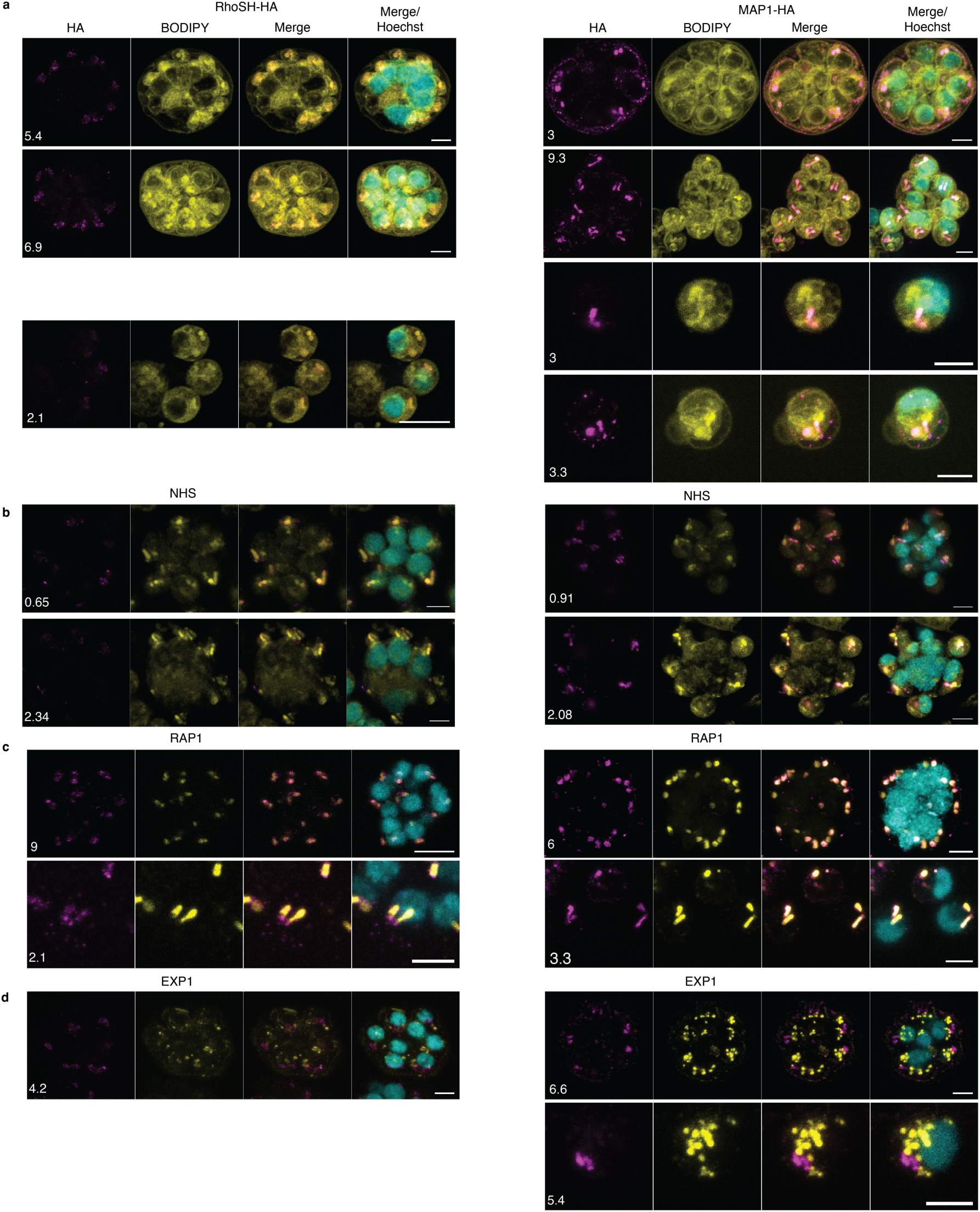
Ultrastructure expansion microscopy reveals that both proteins localise to the rhoptries. The localisation of RhoSH and MAP1 was determined during the schizont stage using ultrastructure expansion microscopy. Rabbit anti-HA was used to probe for target proteins (magenta), **a** BODIPY TR Ceramide (yellow) for membranes, **b** N-hydroxysuccinimide (NHS) ester (yellow) for protein dense regions, **c** RAP1 (yellow) for rhoptry bulb, d EXP1 (yellow) for dense granules. Hoechst was used to stain the nucleus. All images are maximum-intensity projections of a z-stack, where the number refers to the thickness of the stack in µm. Scale bar = 5 µm.

### 3. Co-immunoprecipitation assays reveal that RhoSH and MAP1 interact with each other

To elucidate the protein interactomes of MAP1 and RhoSH, we performed co-immunoprecipitation assays in schizont-stage parasites, followed by mass spectrometry, using WT parasites as a negative control. Percoll-purified parasite pellets containing late-stage schizonts were either saponin lysed and treated with 1% Triton X-100 or lysed directly in 2% digitonin. For the Triton X-100 condition, not many proteins were significantly co-precipitated with either RhoSH or MAP1 when compared to the WT control **(Fig. S4)**. Interestingly, RhoSH co-precipitated three reticulocyte-binding-like (RBL/Rh) homologues (PBANKA_0501000, PBANKA_0316600, and PBANKA_0216801) as well as a merozoite surface protein (PBANKA_0519100), although only one of the RBLs that co-precipitated with RhoSH (PBANKA_0501000) was significantly enriched relative to the WT control **(Fig. S4a)**. MAP1 co-precipitated the Rhoptry-associated, leucine zipper-like protein 1 (RALP1, PBANKA_0619700), which localises to the rhoptry neck (Haase et al., 2008; Ito et al., 2013) and exhibits partial membrane association (Haase et al., 2008) **(Fig S4b)**.

We additionally performed immunoprecipitation assays on digitonin lysed parasite pellets. Interestingly, during this condition, MAP1 and RhoSH reciprocally co-precipitated, indicating a direct interaction. Both proteins were also associated with PBANKA_1312200, a putative ubiquitin-protein ligase whose *P. falciparum* orthologue (PF3D7_1448400) has been linked to potential artemisinin resistance (Cerqueira et al., 2017; Rosenthal & Ng, 2020), as well as PBANKA_0614000 **(Fig. 3a)**. RhoSH-3xHA co-precipitated PIR and fam-b, two proteins exported into the RBC (Fougère et al., 2016). MAP1-3xHA was associated with several exported and rhoptry proteins, including the T-cell immunomodulatory protein (TIP) (Kalia et al., 2021), the exported protein PBANKA_0100700 (Marti et al., 2004), and, like in the Triton-X 100 condition, RALP1, further supporting the relationship of these two rhoptry proteins **(Fig. 3a).** It is possible that 1% Triton-X 100 disrupted the interaction of RhoSH and MAP1, given that digitonin can preserve protein complexes more effectively compared to Triton-X 100.

**Figure 3:**
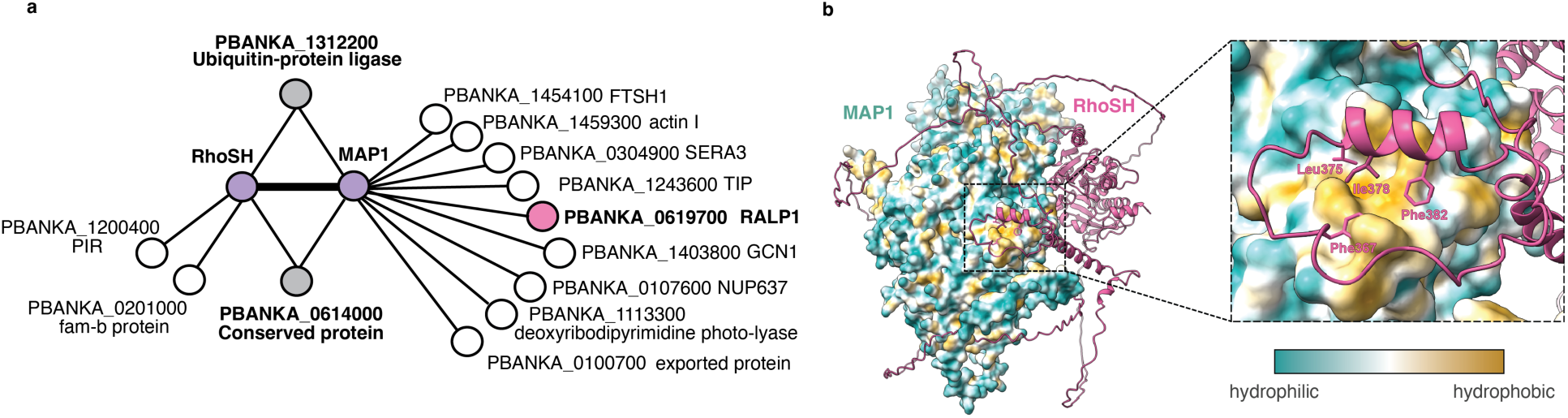
Co-immunoprecipitation assays and AlphaFold predictions indicate that RhoSH and MAP1 are interacting with each other during the schizont stage. a Pellets enriched with schizont stage parasites were lysed in digitonin, and the lysate incubated with anti-HA agarose beads and bead eluate analysed using mass spectrometry. Both RhoSH and MAP1 (in purple) co-precipitated each other and also co-precipitated two other proteins (in grey), PBANKA_1312200 and PBANKA_0614000. MAP1 also co-precipitated a rhoptry neck protein, RALP1 (in pink). Proposed interaction network only shows proteins with significant interaction compared to wildtype control from three biological replicates. b AlphaFold2 model of the RhoSH-MAP1 complex (left) with a zoom-in panel (right) of the predicted interaction site. MAP1 is visualised as a molecular surface, coloured according to hydrophobicity (calculated in ChimeraX), and the RhoSH protein model is indicated in pink. The zoom-in panel shows the predicted interaction site with MAP1 forming a hydrophobic pocket near the hydrophobic peptide side chains of RhoSH. The colour key indicates hydrophilic and hydrophobic regions of MAP1.

To explore potential structural interactions, we used AlphaFold2 multimer (Evans et al., 2021) to model the RhoSH-MAP1 complex. The predicted structure suggests a hydrophobic interface between the two proteins, consistent across top-scoring models despite overall low confidence. In these predictions, the interaction is mediated by RhoSH residues Phe367, Ile378, Phe382 and Leu375, which interact with a hydrophobic pocket in MAP1 (residues Tyr435, Ile438, Ile439, Ile442, Phe446, Leu450, Leu453) **(Fig. 3b, S1c-d)**. In contrast, AlphaFold2 multimer did not predict direct interactions between RhoSH or MAP1 with either PBANKA_1312200 or PBANKA_0614000, suggesting these associations may be indirect. Together, these findings reveal a direct MAP1-RhoSH interaction, along with associations with exported proteins and potential virulence factors.

### 4. Δ*rhosh* and Δ*map1 P. berghei* parasites display reduced pathogenicity and a sequestration defect *in vivo*

To explore the roles of RhoSH and MAP1 in malaria pathogenesis, we generated knockout (KO, indicated with Δ) lines using *Plasmo*GEM vectors (Bushell et al., 2017; Schwach et al., 2015) in the PbmCherry*_hsp70_*+Luc*_eef1_*_α_ (mCherry-Luc) background line. This parental line constitutively expresses mCherry (under the *hsp70* promoter) and firefly luciferase (under the *eef1α* promoter) (Prado et al., 2015). Successful generation of transgenic lines was confirmed by genotyping PCRs **(Fig. S2a,c)**, and clones were obtained by limiting dilution before experiments.

Mice were infected with either Δ*rhosh* or Δ*map1* parasites, and parasitemia, weight, and clinical symptoms were monitored daily **(Fig. 4a)**. Mice infected with the parental mCherry-Luc line served as controls. Both KO lines showed significantly increased survival compared to control-infected mice, which consistently developed moderate symptoms and required euthanasia around day 8 (median) **(Fig. 4bi, c)**. Notably, all Δ*map1*-infected mice (n = 12) survived through the entire experiment (28 days), showing only mild early symptoms and subsequently recovering and gaining weight **(Fig. 4bi, ciii, S5a)**. Δ*rhosh*-infected mice exhibited reduced virulence, with a median survival of 21 days, at which point they were euthanised upon exhibiting moderate clinical symptoms such as hunching and piloerection, accompanied by weight loss **(Fig. 4bi, cii, S5a)**. Daily parasitemia counts revealed a reduced growth rate for both KO lines compared to the mCherry-Luc control during early stages of infection **(Fig. 4bii-iii)**. For Δ*rhosh*, parasitemia rose sharply from day 10 onward, which correlated with the symptom development, and reached 60% at the endpoint, correlating with the onset of moderate symptoms **(Fig. 4b-c)**. In contrast, Δ*map1* parasites displayed a transient increase in parasitemia followed by a decline, suggesting effective host immune control of infection **(Fig. 4bii-iii)**.

**Figure 4:**
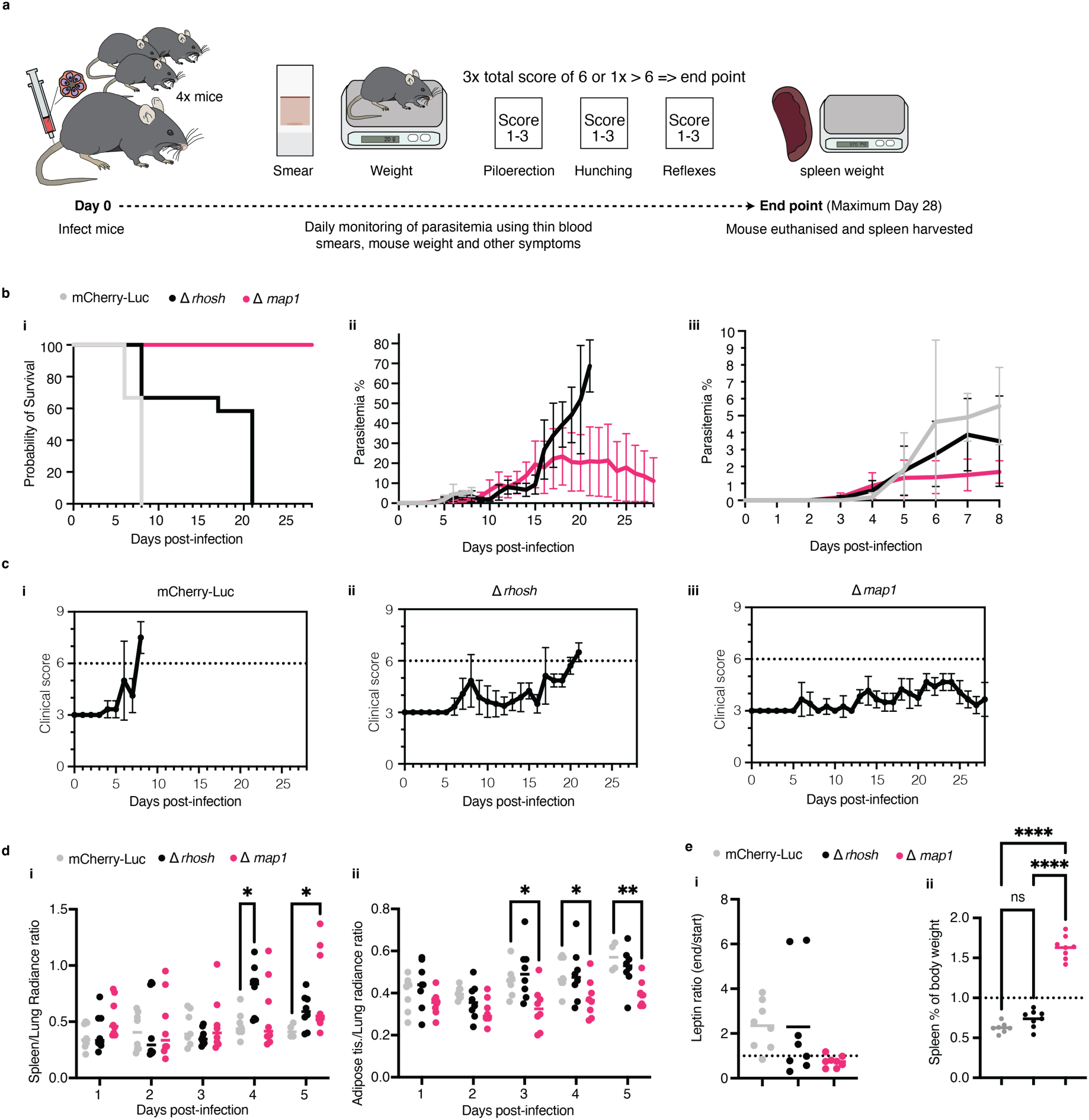
Both Δ*rhosh* and Δ*map1* parasites display reduced virulence and sequestration *in vivo*. **a** Schematic overview of the pathology experiments to determine parasite virulence *in vivo*. For each experiment, four mice were infected and monitored on a daily basis over a period of 28 days (experimental time frame), using thin blood smears, weight measurements and clinical scoring of malaria symptoms (piloerection, hunching, reflexes). If a mouse achieved a combined score equal to six during three consecutive monitoring occasions, or a score above six, the mouse was immediately euthanised and spleen harvested for weighing. **b i** Mice were infected with Δ*rhosh*, Δ*map1* or mCherry-Luc parasites and monitored over 28 days (see panel a) and the probability of survival was plotted., **ii** Giemsa-stained smears from tail-bleed were used to monitor parasitemia. **iii** A zoomed in part (day 0 to 8) of the graph displayed in **ii**. Graphs display data from three individual experiments, where each experiment included four mice per condition. **c** Clinical scores from all pathology experiments shown in panel (b), for **i** mCherry-Luc, **ii** Δ*rhosh* and **iii** Δ*map1*. The data collected for mCherry_Luc control presented in panel (b) and (c) was also included in Hernandez et al., 2024, as these experiments were completed at the same time to reduce the use of mice. **d** *In vivo* imaging system (IVIS) was used to determine parasite sequestration *in vivo*, where mice infected with Δ*rhosh*, Δ*map1* or mCherry-Luc were injected with D-luciferin and the luminescence of luciferase (proxy for parasite density) was measured via imaging. Luciferase signal in **i** the spleen with P-values for Δ*rhosh* (p = 0.0183) and Δ*map1* (p = 0.0299) and **ii** adipose tissue with P-values for Δ*map1* (day 3, p = 0.0126; day 4, p = 0.0262; day 5, p = 0.0041), where mutant lines were compared to the mCherry-Luc control parasites. Statistical analysis was done using two-way ANOVA from two individual experiments, where each experiment included four mice per condition. Images for each timepoint are shown in **Fig. S6**. The data collected for biological replicate one for the mCherry_Luc control was also included in (Hernandez et al., 2024), as these experiments were completed at the same time to reduce the use of mice. **e i** Baseline leptin levels were taken from mice prior to infection with Δ*rhosh*, Δ*map1* or mCherry-Luc parasites. Serum was collected on day eight post-infection, to measure end-point leptin levels and **ii** spleen was harvested and weighed and compared to body weight. Statistical analysis was done using two-way ANOVA from two individual experiments, where each experiment included four mice per condition.

We used the *in vivo* imaging system (IVIS) to assess iRBC sequestration in the luciferase-expressing KO lines. Compared to mCherry-Luc control mice, an increased spleen signal was observed for Δ*rhosh* at day four (p = 0.0183) and for Δ*map1* at day five (p = 0.0299), indicating a potential impaired sequestration resulting in increased splenic clearance for both KO mutants. Importantly, only Δ*map1* displayed a significantly reduced adipose tissue signal starting on day three, consistent with a sequestration defect **(Fig. 4d, S5b)**. These results suggest that the strongly reduced virulence observed for Δ*map1* is at least partly due to failure of iRBCs to sequester to adipose tissue. The sequestration of iRBC to adipose tissue is known to increase secretion of the adipokine leptin by adipocytes, resulting in elevated systemic leptin levels, which correlate with severe malaria in humans and in the *P. berghei in vivo* model (Mejia et al., 2021). We therefore measured systemic leptin levels at day zero and eight post-infection. Infections with the mCherry-Luc control line and Δ*rhosh* parasites yielded a ∼2-fold increase in leptin by day eight, whereas leptin levels remained unchanged in Δ*map1*-infected mice (**Fig. 4ei**). Consistently, Δ*map1*-infected mice exhibited significantly increased spleen-to-body ratios, indicating enhanced splenic clearance, likely linked to reduced sequestration **(Fig. 4eii)**.

Taken together, both RhoSH and MAP1 influence disease progression *in vivo*. Δ*rhosh* parasites exhibit reduced virulence, with delayed symptom onset and increased survival compared to control parasites, while Δ*map1* parasites cause a non-lethal infection and an absence of moderate symptoms during the experimental timeframe. Both KOs show increased parasite loads in the spleen. However, only Δ*map1* exhibits impaired adipose tissue sequestration. This sequestration defect is associated with reduced systemic leptin levels, which likely contribute to the protection of Δ*map1*-infected mice from severe disease **(Fig. 4d-e)** (Mejia et al., 2021).

### 5. Both Δ*rhosh* and Δ*map1* parasites exhibit PV/PVM defects and abnormalities in rhoptry morphology

Since both RhoSH and MAP1 were localised to the rhoptries in this study, we decided to examine late schizont stage development and specifically look at rhoptry morphology using U-ExM. NHS-ester was used to stain for the rhoptries and RAP1 for the rhoptry bulb. Overall, the rhoptries in Δ*rhosh* and Δ*map1* schizonts looked more irregular compared to mCherry_Luc schizonts, and often clustered together **(Fig. 5a)**. When looking specifically at rhoptries in individual merozoites, Δ*map1* had significantly more abnormal rhoptries when compared to mCherry_Luc **(Fig. 5b)**. This included merozoites that had one or more than two rhoptries, a strange shape, or fragmentation/multiple small membranes that contained RAP1 and NHS-ester.

**Figure 5:**
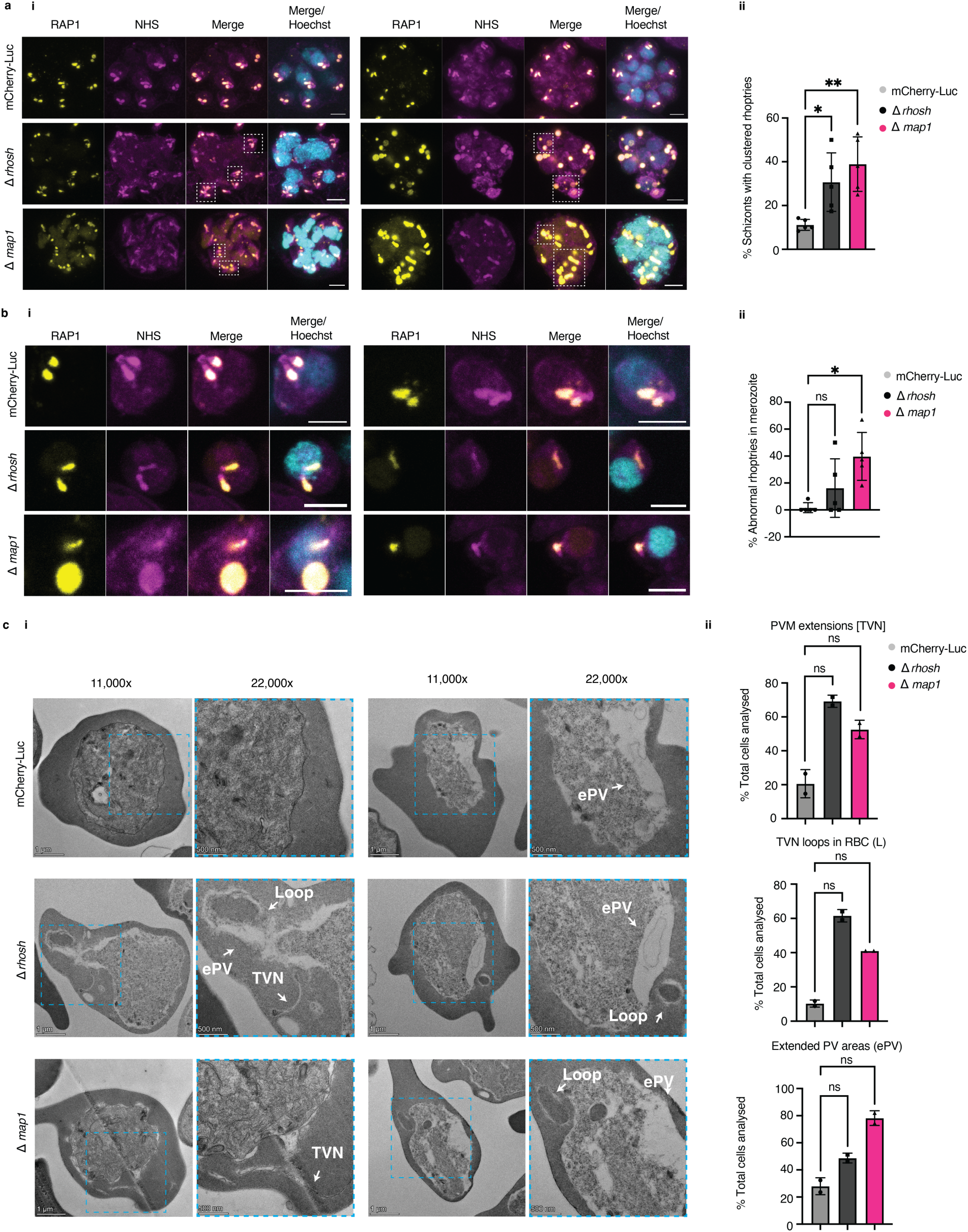
Both Δ*rhosh* and Δ*map1* parasites display abnormal rhoptry and PV/PVM morphology. **a** Ultrastructure expansion microscopy was used to analyse late schizont stage mCherry_Luc, Δ*rhosh* and Δ*map1* parasites. **i** Example of images analysed from five biological replicates, where a white dotted box is used to indicate irregular rhoptry clusters. **ii** Both Δ*rhosh* and Δ*map1* parasites had significantly more (p = 0.0466 and p = 0.00466, Kruskal Wallis test) irregular clusters of rhoptries compared to the mCherry_Luc control. Error bars represent standard deviation for the five biological replicates analysed per condition. **b** Individual merozoites analysed from the same biological replicates presented in panel (a). **i** Examples of individual merozoites analysed. **ii** Merozoites were considered to have abnormal rhoptries if the number of rhoptries was either one or more than two. Only Δ*map1* parasites had significantly different rhoptry populations compared to mCherry_Luc parasites (p = 0.011, Kruskal Wallis test). Error bars represent standard deviation for the five biological replicates analysed per condition. **c** Transmission electron microscopy analysis of trophozoite stage mCherry_Luc, Δ*rhosh* and Δ*map1* parasites. **i** Examples of individual cells analysed, where relevant morphological changes are labelled with white arrows in a zoomed-in version indicated with a blue dotted box. **ii** Graphs show results from differences observed in PVM extensions or the tubulovesicular network (TVN), TVN loops within the iRBC (L), and expansion of the PV (ePV). Error bars represent standard deviation for two individual counters, where two biological replicates were grouped together per condition. Scale bars = 1 μm in standard image and scale bars = 500 nm in zoomed in version.

To further characterise parasite morphology in Δ*rhosh* and Δ*map1* parasites, we used transmission electron microscopy (TEM) to examine membrane morphology during the trophozoite stage. Both Δ*rhosh* and Δ*map1* showed more membrane extensions from the PV, likely a proliferation of the tubulovesicular network (TVN) (Matz, Goosmann, et al., 2015) **(Fig. 5c)**. We also observed more vacuole-like structures inside the RBC, which we referred to as “TVN loops” as they were sometimes connected to the TVN structures, similar to what has been described for depletion of EXP1 (Nessel et al., 2020) **(Fig. 5c)**. Additionally, both Δ*rhosh* and Δ*map1* showed areas of what appeared to be expansion of the PV (Sachanonta et al., 2011) **(Fig. 5c)**. TVN abnormalities were more prevalent in Δ*rhosh* parasites and expansion of the PV in Δ*map1* parasites. Although there was an obvious trend observed for Δ*rhosh* and Δ*map1* compared to mCherry-Luc control, this was not significant, possibly due to the restriction of sample size using TEM.

### 6. MAP1 and RhoSH play a role in parasite lipid metabolism and homeostasis

Next, we looked further into the potential mechanism of action of these two proteins. The functional predictions of RhoSH and MAP1, combined with their localisation and KO phenotypes, lead us to hypothesise a potential role of these proteins in PVM membrane biogenesis and lipid metabolism. We therefore performed untargeted mass spectrometry-based lipidomic analysis of Δ*rhosh* and Δ*map1* mutant trophozoites (cultured *ex vivo* for seven hours). To identify any potential effect of the loss of *map1* or *rhosh* on lipid metabolism we determined the lipidomic profile and content of the iRBC, based on their headgroups and the number of molecular species for each lipid class (**Fig. 6a**).

**Figure 6:**
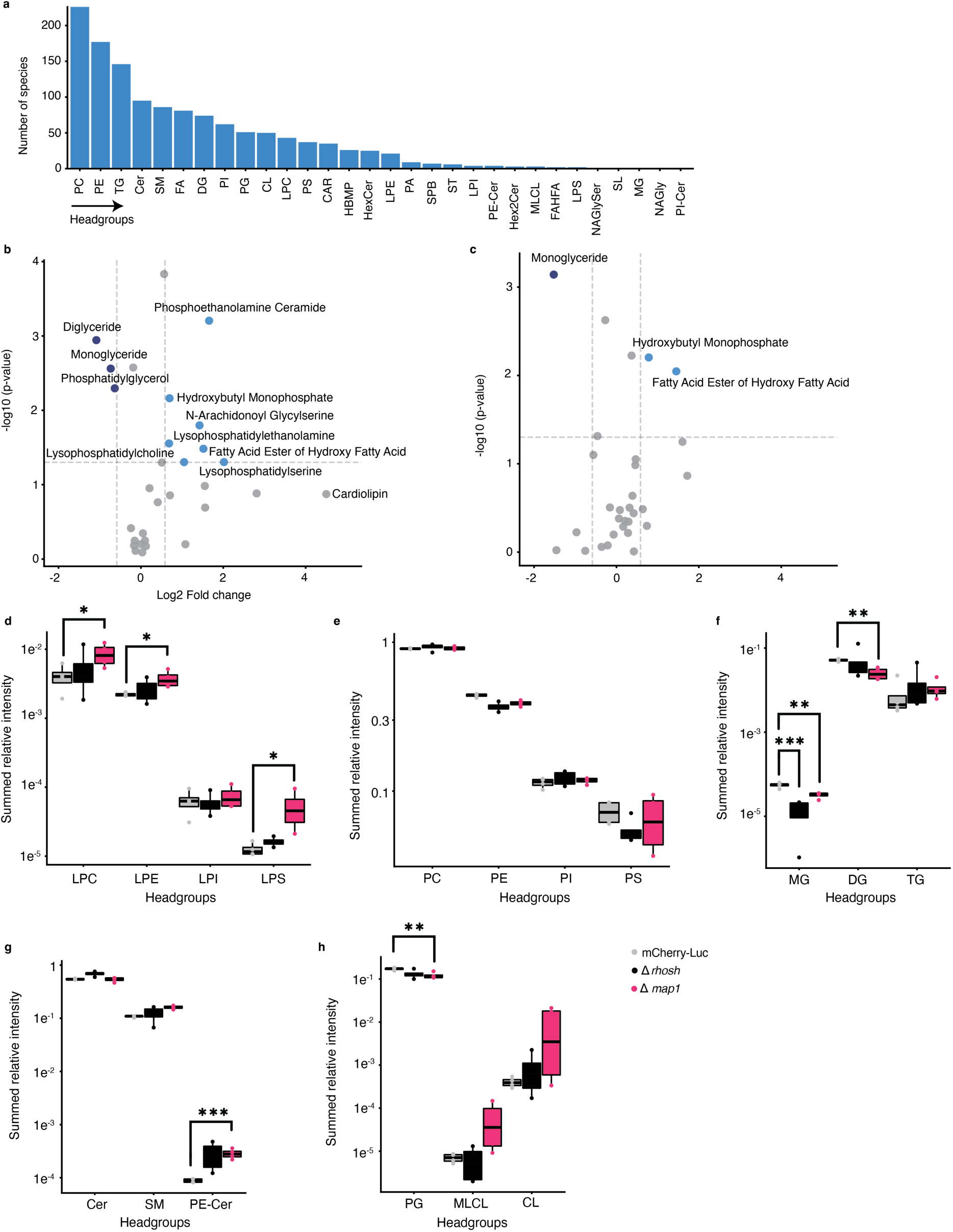
*P. berghei* lipid metabolism is affected by the absence of *map1* and *rhosh*. **a** Grouping of all detected lipid species based on headgroup and ranked in descending order along the x-axis based on the number of different lipid species for each headgroup. Shown in a are only those lipid species detected for all three lines (Δ*map1*, Δ*rhosh* and mCherry-Luc background line), the number of species is therefore identical for all the parasite lines. The intensity of each lipid species was then normalised to the total lipid intensity within that sample and used to calculate the relative abundance of individual lipid species, referred to as summed relative intensity: **b-c** Volcano plots where the Log2 fold change (cutoff 1.5-fold) in lipid abundance between *P. berghei* loss-of-function mutants and the mCherry-Luc background line is plotted against significance (-Log10, p<0.05) for Δ*map1* (a) and Δ*rhosh* (b) mutant lines. **d-h** Boxplots displaying the summed relative intensity of all species within each headgroup for the mCherry-Luc control (grey), Δrhosh (black) and Δmap1 (pink) for selected lysophospholipids (d), glycerophospholipids (e), neutral lipids (f), sphingolipids (g), and cardiolipins. Statistically significant differences in lipid abundances in Δmap1: LysoPC p=0,0497 and LysoPE p = 0,0278, LysoPS p = 0,0496 (d), MG p = 0,0027 and DG p = 0,0011 (f), PE-CER p = 0,0006 (g) and PG p = 0,0050(h), as well as Δrhosh: MG p = 0,0007 (f). Analysis was completed on four biological replicates. Abbreviations: phosphatidic acid (PA), phosphatidylglycerol (PG), phosphatidylcholine (PC), phosphatidylethanolamine (PE), phosphatidylserine (PS), phosphatidylinositol (PI), lysophosphatidylcholine (LPC), lysophosphatidylethanolamine (LPE), lysophosphatidylserine (LPS), lysophosphatidylinositol (LPI), ceramide (Cer), hexosylceramide (HexCer), dihexosylceramide (Hex2Cer), phosphoethanolamine ceramide (PE-Cer), phosphatidylinositol-ceramide (PI-Cer), sphingomyelin (SM), sterol lipids (SL), saccharolipids (SL), sphingoid bases (SPB), fatty acid (FA), cardiolipin (CL), monolysocardiolipin (MLCL), monoglyceride (MG), diglyceride (DG), triglyceride (TG), hemibismonoacylglycerophosphate (HBMP), fatty acyl esters of hydroxy fatty acids (FAHFA), N-acyl glycyl serine (NAGlySer), N-acyl glycyl (NAGly), Acylcarnitines (CAR).

To determine the changes in lipid homeostasis upon the disruption of *map1* and *rhosh*, we measured the most significant changes in lipid molecular composition by quantifying significant fold changes from relative abundances of each lipid for both the *Δmap1* and *Δrhosh* mutant lines (**Fig. 6b-c**). The loss of *map1* but not *rhosh*, induced a significant increase of the content of lysophospholipid classes, most importantly lysophosphatidylethanolamine (LPE), lysophosphatidylserine (LPS) and lysophosphatidylcholine (LPC) (**Fig. 6b-d**). Although milder than in the *Δmap1* mutant, the disruption of *rhosh* also induced a slight, but not significant, increase of LPE, LPC and LPS (**Fig. 6d**). There was no significant change in other glycerophospholipid content (PE, PC or PS glycerophospholipids), in either of the mutants (**Fig. 6e**). However, the loss of either *map1* or *rhosh* resulted in a significant decrease in the neutral lipid monoacylglyceride (MAG) and a significant decrease of a second neutral lipid, diacylglyceride (DAG) in the *Δmap1* mutant (**Fig. 6b-c, f**), together suggesting a potential role of the enzyme to provide fatty acid substrates, putatively from lysophospholipids for the synthesis of neutral lipids. Phosphatidic acid (PA) and DAG together constitute an important nexus in lipid metabolism, acting as precursors for synthesis of both membrane phospholipids and the storage lipid triacylglyceride (TAG), (Carman & Han, 2009).

Next, we investigated the effect of knocking out *map1* or *rhosh* on ceramide and sphingolipid metabolism. No significant differences were seen in ceramide and sphingomyelin levels, although ceramide levels were slightly increased in *Δrhosh* and sphingomyelin levels elevated in *Δmap1*. However, the loss of *map1* resulted in a significant increase of phosphoethanolamine ceramide (PE-Cer), with *Δrhosh* again exhibiting an intermediate, but not significant, increase (**Fig. 6b-c, g**). PE-Cer is produced from ceramide and PE, the latter which donates its polar headgroup in a reaction that results in the release of DAG (Panevska et al., 2019), again linking polar and neutral lipid metabolism. The significant increase in PE-Cer in absence of a clear effect on ceramide or sphingomyelin could indicate that PE-Cer is also serving as a substrate for MAP1 (and to a lesser extent RhoSH) hydrolysis. The only glycerophospholipid that was significantly reduced in the *Δmap1* mutant was phosphatidylglycerol (PG) (**Fig. 6h**). Notably, PG is a precursor for cardiolipin (CL) synthesis. CL is the major structural lipid of mitochondrial membrane and is essential for parasite development and virulence (Fu et al., 2018; Pietsch et al., 2023). Interestingly, we measured an increase of both CL and monolysocardiolipin (MLCL), an intermediate of CL synthesis and degradation. However, the high variability between samples resulted in a lack of statistical significance (**Fig. 6h).**

Taken together, this analysis shows that deletion of *map1*, and to a lesser extent *rhosh*, affects parasite lipid content and homeostasis. The significant decrease in neutral lipid classes MAG and DAG, coupled with the increase in lysophospholipids, could suggest a putative lysophospholipase activity by MAP1, and indirectly by RhoSH. Lipases are characterised by a conserved Gly-X-Ser-X-Gly motif also present in *Plasmodium* lipases and centred on the serine of the active site (Flammersfeld et al, 2018). RhoSH, which contains a canonical Ser125-His218-Asp189 catalytic triad, also has a Gly-Arg-Ser-Leu-Gly motif, while no such Gly-X-Ser-X-Gly motif could be identified in MAP1.

### 7. Characterisation of the MAP1 orthologue in *P. falciparum* supports a role in CD36-mediated cytoadhesion

Given the critical role of MAP1 in *P. berghei* virulence, we next investigated whether its function is conserved in the major human malaria pathogen *P. falciparum* by characterising its orthologue PF3D7_0811600 (PfMAP1). Since this gene was reported essential in an insertional mutagenesis screen (Zhang et al., 2018), we generated a conditional knockdown line by appending to the endogenous locus a 3xHA tag, Neomycin resistance gene (NeoR), and a *glmS* riboswitch, using the selection-linked integration system (Birnbaum et al., 2017; Jonsdottir et al., 2021; Spielmann et al., 2017). The *glmS* system enables conditional downregulation of gene expression upon glucosamine (GlcN) addition (Prommana et al., 2013).

Correct integration of HA-NeoR-glmS at the *pfmap1* locus was confirmed by genotyping PCRs **(Fig. S7)**, and knockdown was confirmed by western blot and IFAs following two cycles (trophozoite to trophozoite) of treatment ± 2.5 mM GlcN **(Fig. 7a-b)**. PfMAP1-HA-NeoR-glmS (referred to as PfMAP1-HAglmS from now on) parasites showed near-complete knockdown (97 and 98% in two independent experiments) **(Fig. 7a)**. The expression of the exported knob-associated histidine-rich protein (KAHRP) and parasite growth were both unaffected after two cycles of GlcN treatment **(Fig. 7b-c)**. Since the growth of the knockdown line was not significantly different from the 3D7 WT control, this suggests that the attenuated phenotype observed in *P. berghei* may be linked to specific host-parasite interactions rather than intrinsic effects on parasite growth or development. Alternatively, residual PfMAP1 levels may suffice to sustain essential protein function. Localisation studies by IFA revealed that PfMAP1-HAglmS co-localises with EXP2 at the PVM (de Koning-Ward et al., 2009) during the early schizont stage but later shifts to the apical end in mature schizonts **(Fig. 7d)**.

**Figure 7:**
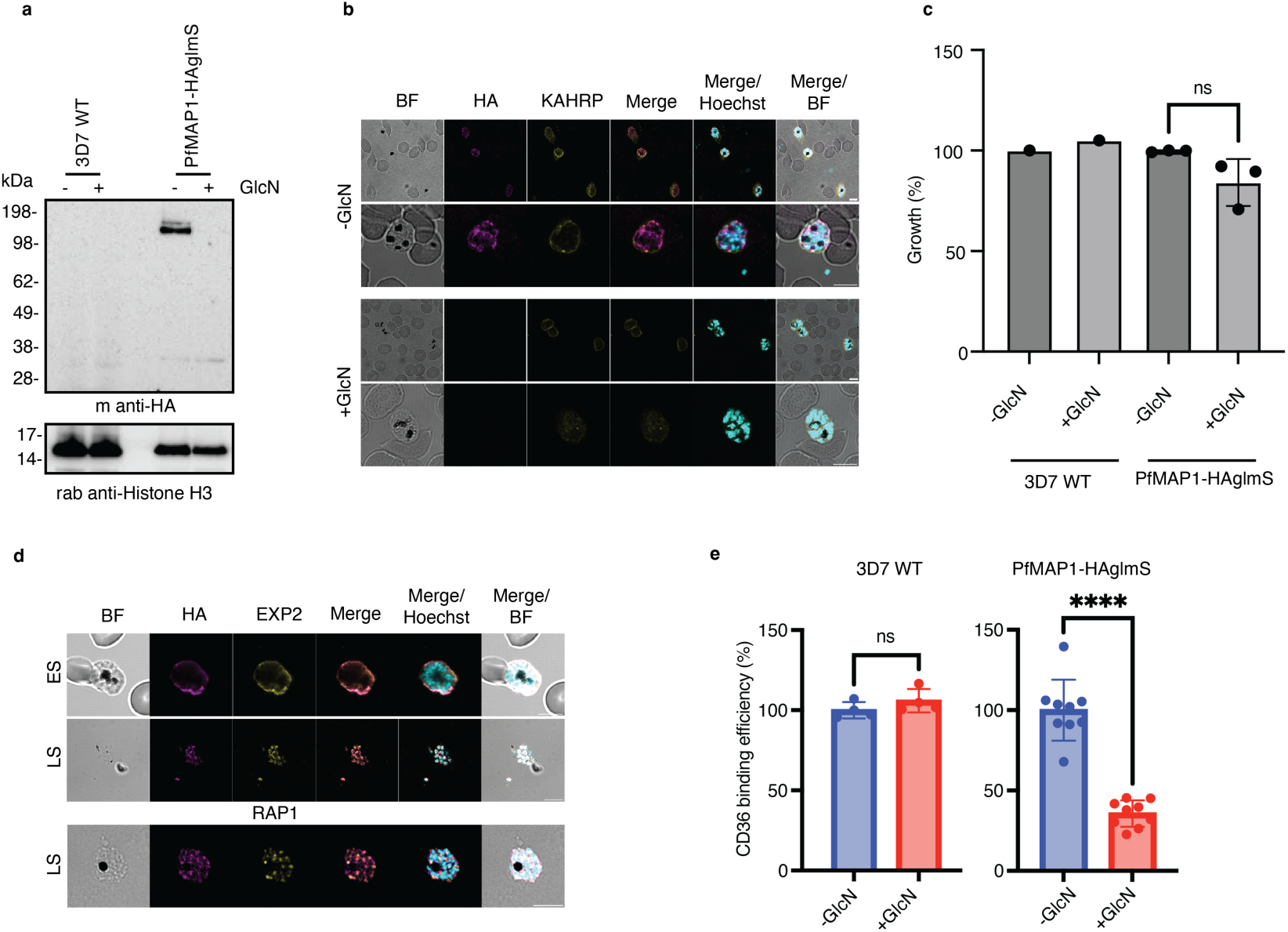
Phenotypic analysis of MAP1 orthologue in *P. falciparum* confirms its role in CD36-mediated sequestration. **a** Western blots prepared from 3D7 WT and PfMAP1-HAglmS schizont stage parasites treated ± 2.5 mM GlcN, probed with mouse anti-HA (tagged protein) and rabbit anti-H3 (loading control). Image is representative of two biological replicates. **b** Immunofluorescence assay on schizont stage parasites treated ± 2.5 mM GlcN for two cell cycles. Rat anti-HA (magenta) was used to probe for the target protein, rabbit anti-KAHRP (yellow) was used as a marker for an exported protein, and Hoechst (cyan) to stain the nucleus. Scale bar = 5 μm. BF = bright field. **c** 3D7 WT and PfMAP1-HAglmS schizont stage parasites were treated ± 2.5 mM GlcN for two cell cycles. Statistical analysis was completed using Student’s t test from three biological replicates, error bars = standard deviation. **d** Immunofluorescence assay completed for PfMAP1-HAglmS on early and late schizont stage (ES and LS), where rat anti-HA (magenta) was used to probe for the target protein, mouse anti-EXP2 (yellow) used as a marker for the PVM and dense granules, mouse anti-RAP1 (yellow) used as marker for the rhoptries, and Hoechst (cyan) to stain the nucleus. Scale bar = 5 μm. BF = bright field. **e** 3D7 WT and PfMAP1-HAglmS parasites were treated ± 2.5 mM GlcN for two cell cycles and binding to CD36 was assessed. Statistical analysis was done using Student’s t test on three biological replicates, where **** represents a P-value <0.0001.

To determine whether the reduced sequestration phenotype observed for Δ*map1* in *P. berghei* is conserved in *P. falciparum*, we assessed PfMAP1 function in a cytoadhesion assay, where control and mutant iRBCs were incubated with purified CD36 that had been immobilised on a solid surface. Knockdown of PfMAP1 resulted in a significant decrease in binding to CD36, the primary receptor mediating sequestration on microvascular endothelial cells, whereas no difference was observed in 3D7 WT parasites treated with GlcN **(Fig. 7e)**. These findings indicate that PfMAP1 likely contributes to sequestration and CD36-mediated cytoadhesion in *P. falciparum*, and that this function is conserved between *P. berghei* and *P. falciparum*.

## Discussion

Our findings identify MAP1 and RhoSH as critical contributors to the pathology of malaria parasites. Both proteins were previously identified to be important for normal asexual parasite growth as part of a large-scale genetic screen (Bushell et al., 2017) and here we provide a detailed characterisation of these two proteins and their importance *in vivo*.

During the *P. berghei* trophozoite stage MAP1 was observed lining the PVM, sometimes on the iRBC side of EXP1, within distinct regions that did not appear to overlap with concentrated areas of EXP1 at the PVM. In both *P. berghei* and *P. falciparum,* MAP1 is then observed in the apical organelles in late segmented schizonts, where we confirm that in *P. berghei* MAP1 is localised to the rhoptry bulb membrane, surrounding RAP1 within the lumen (Liffner et al., 2020; Richard et al., 2009). RhoSH was sometimes observed in the proximity of the PV/PVM during the early schizont stage, and did not appear to concentrate in the same regions as EXP1. RhoSH was apically localised during later schizogony, forming distinct puncta surrounding the rhoptries, and often concentrated around the rhoptry neck. Full-length MAP1 was partially soluble, as was both the processed and some of the full-length form of RhoSH. Both MAP1 and RhoSH were mostly peripherally associated with membranes, where the majority of the full-length form of RhoSH was released with carbonate buffer, whilst the processed form was similarly divided between soluble and peripheral membrane association. This suggests that the full-length form of RhoSH has a stronger membrane association, which may help explain the difference in its localisation compared to MAP1 within/around the rhoptries. Prodomains of rhoptry proteins have been shown to mediate trafficking, enhance stability and secretion, or maintain zymogen forms until activation (Counihan et al., 2013). In *P. falciparum,* RhoSH has been identified as an active insoluble α/β serine hydrolase in proteomic studies (Davison et al., 2022; Elahi et al., 2019), suggesting potential as a druggable enzyme. In addition, the activity of RhoSH was observed to increase as the parasite develops from schizonts to merozoites (Davison et al., 2022), supporting activation of RhoSH within the secretory organelles upon merozoite egress.

Interestingly, RhoSH and MAP1 co-precipitated during late schizogony, when both proteins are observed around the rhoptries, suggesting a possible shared function at this stage and potentially at the PV/PVM where they both localise to during the early stages of schizogony. This interaction was further supported by AlphaFold modelling. Additionally, MAP1 co-precipitated RALP1, a known rhoptry neck protein, which is also partially associated with membranes but lacks a predicted transmembrane domain (Haase et al., 2008). Although antibodies against PfRALP1 inhibit parasite invasion (Ito et al., 2013), we saw no evidence of an invasion defect in PfMAP1 knockdown parasites, suggesting that MAP1 and RALP1 might not have shared function during invasion, although a complete PfMAP1 knockout is needed to confirm this.

Both Δ*rhosh* and Δ*map1 P. berghei* parasites showed reduced virulence *in vivo*, where mice infected with Δ*map1* failed to develop moderate symptoms throughout the duration of the experiment. During the early stages of infection, both mutants grew slower compared to the control, and parasitaemia started to rise around day five of infection. For Δ*map1* infected mice, the parasitemia plateaued around day 15, whilst the parasitemia in Δ*rhosh* mice showed exponential growth from day 15 until the mice were euthanised on day 21. Both mutant lines showed increased parasite accumulation in the spleen, consistent with impaired cytoadhesion. Δ*map1* parasites also showed significantly reduced sequestration to adipose tissue, further supporting defects in cytoadhesion. The impaired sequestration coupled with reduced leptin levels and initial lower parasitemia likely explains the strongly reduced virulence of Δ*map1* parasites. In line with our findings and during the course of this study, Gao et al., 2023 independently reported reduced virulence in mice infected with Δ*map1* parasites. Their study showed extended host survival but not full protection, likely reflecting differences in inoculum size. Interestingly, they demonstrated that immunisation with MAP1-derived peptides significantly prolonged survival, and sera from these mice conferred passive immunity. Moreover, Δ*map1* parasites elicited stronger IFN-γ and TNF-α responses than WT parasites, suggesting that MAP1 expression suppresses protective immunity and underscoring its potential as a recombinant vaccine candidate (Gao et al., 2023).

Significantly, knockdown of PfMAP1 resulted in reduced binding of iRBCs to the endothelial receptor CD36 *in vitro*, strongly indicating a conserved role of MAP1 in *P. falciparum* and further supporting that MAP1 may play a role in immune evasion through sequestration and reduced splenic clearance *in vivo*. However, export of KAHRP did not seem to be affected when PfMAP1 was knocked down, indicating that protein export is not globally disrupted. This pattern is consistent with previous observations for PTEX88 and PV1 (both localised in the PV), where knockout or knockdown parasites displayed impaired sequestration, but their function could not directly be linked to protein export (Chisholm et al., 2016; Matz, Ingmundson, et al., 2015).

When assessing the effect of losing either *map1* or *rhosh* on PVM morphology, we observed increased prevalence of areas of PVM expansion, where the PVM appeared separated from the parasite plasma membrane, as well as increased formation of TVN extensions and vacuole-like structures (loops) from the PVM within the iRBC cytoplasm. These membrane abnormalities of the PVM could potentially have a knock-on effect on the trafficking of cytoadhesive proteins across the PVM, although not completely disrupting export since the mutant parasites are viable and the export of KAHRP remains unaffected. This model is consistent with the suggestion by Davison et al., 2022 that conditional mutation of the catalytic triad in *P. falciparum rhosh* disrupts PVM formation, although experimental evidence was not provided. Additionally, when looking at late-stage schizonts, we observed that a subset of schizonts displayed abnormal rhoptry morphology and altered rhoptry numbers, with more noticeable defects observed in Δ*map1* compared to Δ*rhosh* parasites. However, this does not appear to markedly affect parasite invasion given their ability to complete the parasite intraerythrocytic cycle.

The establishment of the PVM is intimately linked to parasite lipid metabolism and homeostasis. Knockout of *map1* resulted in a significant increase in LPC, LPE, and LPS, and a significant decrease in the neutral storage lipids MAG and DAG. The effect on lipid metabolism upon loss of *rhosh* was less pronounced, with only MAG significantly reduced. Nevertheless, Δ*rhosh* mutant parasites overall displayed an intermediate phenotype to that of Δ*map1*. Hydrolysis of glycerophospholipids results in the release of a free fatty acid and their respective lysophospholipids, which can in turn be hydrolysed to release its polar head and remaining fatty acid. An integral part of parasite lipid metabolism is the careful management of excess free fatty acids, which, to prevent lipotoxicity, are converted via DAG into TAG and stored in lipid droplets (Lee et al., 2024). The conversion of phospholipids into lysophospholipids is catalysed by phospholipases, encoded by an extended repertoire of phospholipase genes in *P. falciparum* (Flammersfeld et al., 2018). In *P. falciparum* the conditional knockdown of lysophospholipase 3 (PfLPL3) results in reduced levels of DAG and TAG (Sheokand et al., 2023), while the knockdown of lysophospholipase 1 (PfLPL1) causes reduced levels of TAG and increased levels of DAG (Asad et al., 2021). Intriguingly, PfLPL3 also localises to the host-parasite interface and is associated with the PV and the TVN (Sheokand et al., 2023). Based on the accumulation of lysophospholipids and reduction in MAG and DAG, we here hypothesise that MAP1, and likely RhoSH, could potentially be lysophospholipases. Work is currently ongoing to identify and experimentally validate their specific substrates by analysis at the level of individual lipid species.

Alongside glycerophospholipids, sphingolipids are important structural membrane components. The knockout of *map1* does not affect the abundance of sphingomyelin, however, the level of PE-Cer is significantly elevated in the absence of MAP1, where again the knockout of *rhosh* results in a milder phenotype. PE-Cer is the dominant sphingolipid of cell membranes in invertebrates, it’s enriched in *Toxoplasma gondii* infected cells but cannot be scavenged from the host by *T. gondii* (Nyonda et al., 2022; Panevska et al., 2019). Significantly, the knockout of *T. gondii* serine palmitoyltransferases 1 (TgSPT1) results in a reduction of PE-Cer, which translates into a rhoptry secretion and invasion impairment phenotype (Nyonda et al., 2022). Whether the effect on PE-Cer and lysophospholipids is linked by MAP1 (and potentially RhoSH) mediating hydrolysis of substrates with specific fatty acid chain length remains to be elucidated. The individual contributions of the different lipid phenotypes to the overall morphological, growth and virulence phenotypes of Δ*map1* and Δ*rhosh* also remain to be determined.

In summary, this study provides a detailed characterisation of two *Plasmodium* virulence factors, RhoSH and MAP1, that modulate parasite growth, sequestration, and pathology. Both proteins putatively act at the PVM as potential lysophospholipases, where they appear to be required for PVM integrity. Their exact lipid substrates and the precise molecular mechanisms by which they potentially influence trafficking of cytoadhesins and thereby affect sequestration remains to be discovered. Nonetheless, their rhoptry-based secretion and the strong mutant phenotypes described here support MAP1 and RhoSH as promising candidates for translational applications. The predicted enzymatic domains of both proteins represent attractive targets for inhibitor design. By linking basic functional insights with translational potential, this study lays the groundwork for further exploration of these proteins as therapeutic and vaccine targets in malaria control efforts.

## Materials and methods

### 1. Protein selection, sequence analysis, and structure predictions

We utilised a spatial proteomics dataset based on density gradient separation of subcellular compartments from *P. falciparum* schizont stages (Chisholm et al., 2025) to identify uncharacterised proteins associated with the PV and its membrane, or clustered with known exported proteins. The selection criteria included (1) confirming the gene had not been characterised before, (2) assessment of gene essentiality for blood-stage development using an insertional mutagenesis screen in *P. falciparum* (Zhang et al., 2018) and (3) cross-referencing with orthologues in *P. berghei* that were identified using VEupathDB resources (OrthoMCL database, release 6.21, (Fischer et al., 2011). The final list of *P. berghei* genes was selected considering the availability of tagging and knockout vectors from the *Plasmo*GEM open access research resource (https://plasmogem.umu.se/pbgem/) (Bushell et al., 2017; Schwach et al., 2015).

The AlphaFold2 models for RhoSH and MAP1, as monomers and heterodimers, were generated with ColabFold (v1.5.2) (Mirdita et al., 2022) using the alphafold2_multimer_v3 model type employing 20 prediction cycles. First, monomeric RhoSH and MAP1 structures were predicted. The best-ranked AlphaFold2 models of MAP1 and RhoSH were used to identify functional domains with Foldseek (van Kempen et al., 2024). The 3Di/AA mode was employed for MAP1, and both 3Di/AA and TM-align modes for RhoSH. For the heterodimer prediction, the best-ranked MAP1 model was used as a template to predict the heterodimeric RhoSH-MAP1 complex. In all cases, the model confidences are indicated by colouring according to the pLDDT (predicted local-distance difference test) confidence measure and the predicted aligned error (PAE) plot **(Fig. S1)** (Jumper et al., 2021).

For signal peptide prediction, we used TargetP 2.0 (Almagro Armenteros et al., 2019), SignalP6.0 (Teufel et al., 2022), Phobius (Käll et al., 2004), DeepTMHMM (Hallgren et al., 2022) and TOPCONS (Tsirigos et al., 2015).

### 2. Routine housing, handling, and sampling of mice

All animal studies were conducted at Umeå University following Ethics Permits A34-2018 and A24-2023, granted by the Swedish Board of Agriculture (Jordbruksverket), and all animals used were purchased from Charles River Europe. A minimum age of six-week-old female BALB/c mice were used for routine parasite infections, and six-week-old female C57BL/6 mice were used for pathology experiments. They were housed in groups of four in individually ventilated cages with autoclaved wood chips and paper towels for nesting, kept at 21 ± 1 °C with a 12-hour light/dark cycle and 55 ± 5% relative humidity. Animals were maintained in specific pathogen-free (SPF) conditions, with Exhaust Air Dust (EAD) monitoring conducted twice a year. All animals had *ad libitum* access to a commercial dry rodent diet and fresh water. Routine daily health monitoring was performed visually. To assess parasitemia, thin blood smears were prepared from tail bleeds, fixed with methanol, and stained with Giemsa 10%. For blood collection, mice underwent submandibular bleeding without anaesthesia, or a terminal bleed under anaesthesia (225-270 mg/kg Ketamine (MSD Animal Health); 47-60 mg/kg Xylazine (Dechra Veterinary Products) in PBS), followed by cardiac puncture using a syringe with or without heparin (50 mg/mL, Sigma-Aldrich). Animals were euthanised by cervical dislocation.

### 3. Generation of *P. berghei* transgenic lines

For generation of *P. berghei* KO and tagged lines, *Plasmo*GEM vectors were prepared and verified by PCR with the vector-specific QCR2 primer and the generic GW2 primer **(Fig. S2a)**. The genotyping primer sequences for all *Plasmo*GEM vectors were obtained from (https://plasmogem.serve.scilifelab.se/pgem/home). Before transfection, the plasmids were linearised with NotI and ethanol precipitated.

Edited *P. berghei* parasite lines were generated in the *P. berghei* ANKA cl15cy1 wild-type strain (tagged lines) or in the PbmCherry*_hsp70_*+Luc*_eef1_*_α_ (mCherry-Luc) background line (KO lines) (Prado et al., 2015). For transfections, schizonts were prepared using infected blood from mice following a modified protocol from Janse et al., 2006. In brief, parasites were cultured for 22 hours in complete medium (RPMI 1640 (Gibco) with 25% fetal bovine serum (Gibco), penicillin-streptomycin (Gibco) at 100 U/mL and 100 µg/mL, respectively, and 24 mM NaHCO3) in gas-sealed flasks (3% CO2, 1% O2, 96% N2) at 37 °C with shaking at 80 rpm. Schizonts were isolated using a 15.2% Histodenz/PBS (Sigma-Aldrich) cushion, washed in complete medium, and transfected with 10 µg of NotI digested *Plasmo*GEM plasmid DNA, by electroporation with the Lonza 4D Nucleofector System (pulse program FI-115) and P3 Primary Cell 4D-Nucleofector X Kit S (Lonza). Post-transfection, parasites were resuspended in 100 µL RPMI and immediately injected intravenously into a mouse’s lateral caudal vein. Selection of resistant transgenic parasites began the day after transfection (day 1) with pyrimethamine (0.07 mg/mL, MP Biomedicals) administered in drinking water.

To evaluate successful genome editing (KO or tagging) by vector integration, PCRs were performed on parasite genomic DNA extracted from 50 μl of infected blood mixed with 150 μl PBS using the DNeasy Blood and Tissue kit (Qiagen) and the gene-specific genotyping GT (integration) primer together with the generic primer GW1 or GW2, depending on the *Plasmo*GEM vector design. Additionally, to detect the WT locus, the gene-specific primers QCR1 and QCR2 were used. The vector-specific QCR2 primer and the generic GW2 primer, which detects both the integrated and unintegrated vector was also used **(Fig. S2a)**. PCR assays were done using GoTaq G2 Green Master Mix (2×, Promega, USA) and cycling conditions included an initial 5 min denaturation at 95 °C, followed by 30 cycles of denaturation at 95 °C for 30 s, annealing for 30 s at 2 °C below the calculated melting temperature of the primers, and elongation at 62 °C for 1 min for every 1 kb, followed by a final extension step of 62 °C for 5 min. All PCR amplicons were analysed using agarose gel electrophoresis stained with Sybr Safe (Invitrogen). The list of all primers and their nucleotide sequence can be found in **Table S1**. To establish *P. berghei* KO clonal lines transfectants were cloned by limiting dilution as described by (Ménard & Janse, 1997).

### 4. Generation of MAP1 knockdown line in *P. falciparum*

To edit *P. falciparum*, we used the selection-linked integration (SLI) method (Birnbaum et al., 2017; Spielmann et al., 2017) using a modified version of the pSLI-3xHA-2A-NeoR-glmS plasmid used in Jonsdottir et al., 2021, where LoxP site had been introduced before the 2A skip peptide and after the NeoR gene, here referred to as pSLI-3xHA-LP-2A-NeoR-glmS **(Fig. S7a)**. The LoxP sites are irrelevant to this study, as plasmids were transfected into 3D7 WT parasites. First, a 918 bp long homology region of the 3’ end of the target gene was PCR amplified using the Cloneamp HiFi PCR premix (Takara) and primers pSLI_Fw_081 and pSLI_Rv_081 (**Table S1**). Cycling conditions included denaturation at 98 °C for 3 min, 25 cycles of denaturation at 98 °C for 10 sec, annealing at 52 °C for 15 sec, and elongation at 62 °C for 30 sec, with a final 5-min incubation at 62 °C. The amplification product was cloned into the modified pSLI vector in the BglII and PstI sites, using Gibson assembly **(Fig. S7a)**. The assembly was transformed into XL10 gold ultracompetent cells (Agilent), and clones were checked by miniprepping and restriction digestion with BglII and PstI. Positive colonies were sequenced with the NeoR_Nterm_5INT_R primer.

Parasite transfections were carried out using 60 μg of plasmid DNA (Macherey Nagel) in cytomix, following standard transfection protocols. Transfectants were initially selected with 2.5 nM WR99210 (WR, Jacobus Pharmaceuticals) to maintain the episomal plasmid. For selection of integrants using the Selection-Linked Integration (SLI) approach, cultures at 2–4% parasitemia were exposed to G418 at a final concentration of 400 μg/mL for neomycin resistance. WR was omitted during this phase. Cultures were fed daily until parasites disappeared (typically within 10 days) and then every other day. Genomic DNA was extracted from emerging parasites using the QIAamp DNA Mini Kit (catalog no. 51304). PCR analyses were performed to confirm integration at the 5’ (PF0811600_5’int_Fw2 and NeoR_Nterm_5INT_R) and 3’ (PF0811600_3’int_Rv1 and pSLI_3INT_F) ends, to assess for the presence of unmodified loci, ruling out WT parasites or incorrect integrations (PF0811600_5’int_Fw2 and PF0811600_3’int_Rv) **(Table S1)**. If traces of the WT locus persisted, cultures were subjected to WR selection to obtain a pure integrant population.

To induce the protein knockdown, sorbitol-synchronised late-stage parasites were cultured for two cycles ± 2.5 mM GlcN (Merck G1514).

### 5. *P. falciparum* routine cell culture

*P. falciparum* strains, including the 3D7 parental line, as well as genetically modified derivatives, were cultured in human erythrocytes obtained from NHS Blood and Transplant (NHSBT, Cambridge, UK). Cultures were maintained at a haematocrit (HCT) of 3-5% in RPMI 1640 medium (Gibco, UK) supplemented with 5 g/L Albumax II, 2 g/L Dextrose Anhydrous EP, 5.96 g/L HEPES, 0.3 g/L Sodium Bicarbonate EP, and 0.05 g/L hypoxanthine, with the latter dissolved in 2 M NaOH. Cultures were maintained in vented-cap flasks or multi-well plates and incubated at 37 °C under a controlled low-oxygen atmosphere (1% O₂, 3% CO₂, 96% N₂; BOC, Guildford, UK) in a gassed incubator.

### 6. Sorbitol synchronisation of *P. falciparum*

Synchronisation of ring-stage parasites was performed following established protocols. Cultures were centrifuged to pellet the cells, and the supernatant was discarded. The pelleted cells were then resuspended in a volume of 5% D-sorbitol (Sigma-Aldrich) equivalent to ten times the pellet size and incubated at 37 °C for 5 minutes. During this step, trophozoites and schizonts were lysed, leaving only ring-stage parasites intact. The treated cells were centrifuged again, washed by resuspension in RPMI medium at twenty times the pellet volume, and pelleted once more. Finally, the cells were resuspended in complete medium and adjusted to a 4% HCT.

### 7. Immunoblot

For *P. berghei*, BALB/c mice were intraperitoneally infected using a frozen stock of parasites. When parasitemia reached ∼5%, blood was collected, and parasites were cultured as previously described. For *P. falciparum*, synchronous cultures (4% HCT) were harvested at late schizont stages. Parasites were enriched either by centrifugation through a 67% Percoll (Cytiva) density gradient (*P. berghei*) or by lysis of infected erythrocytes in 0.1% (w/v) saponin in PBS (*P. falciparum*). In both cases, pellets were washed twice in PBS supplemented with protease inhibitors (cOmplete, Roche) and stored at −80 °C until further use. Parasite pellets were resuspended in sample buffer (1× Laemmli buffer, Bio-Rad, for *P. berghei*; 1× LDS buffer, Invitrogen NP0007, for *P. falciparum*) at a ratio of 20× pellet volume, supplemented with DTT (100 mM for *P. berghei*, 50 mM for *P. falciparum*). Samples were incubated for 5–10 min at 80–90 °C and loaded onto SDS-PAGE gels (4–20% TGX, Bio-Rad, or 4–12% Bolt™ Bis-Tris Plus, Invitrogen). Electrophoresis was carried out at 150 V for 50 min in the corresponding running buffer (Tris-Glycine or MES, respectively). Proteins were transferred onto PVDF (*P. berghei*) or nitrocellulose (*P. falciparum*) membranes (Bio-Rad), pre-soaked in methanol, using a semi-dry transfer system (Bio-Rad) set to 20 V for 7 min. Membranes were blocked for 1 h at room temperature in 1% casein in PBS (*P. berghei*) or 5% skim milk in TBS-T (*P. falciparum*), and then incubated with primary antibodies diluted in blocking buffer, either overnight at 4 °C or for 1 h at room temperature (PfHSP70). After three washes (PBS or TBS-T), membranes were incubated with HRP-conjugated secondary antibodies for 1 h, washed again, and developed using a chemiluminescent substrate (Millipore or Thermo Scientific). Images were captured using a ChemiDoc system (Bio-Rad). Primary antibodies: rat anti-HA (Roche #11867431001, 1:1,000); rabbit anti-Histone H3 (Abcam #ab1791, 1:2,500), rabbit anti-HA (Cell Signaling #C29F4, 1:1,000); rat anti-PbEXP1 (Proteogenix, 1:1,000); rabbit anti-PfHSP70 (LS Bio #LS-C109068-25, 1:10,000); mouse anti-α-spectrin (Abcam #ab11751, 1:1,000). Secondary antibodies: anti-rabbit IgG (H+L) HRP (Jackson ImmunoResearch #111-035-003, 1:10,000; Promega #W4011, 1:2,500), anti-rat IgG (H+L) HRP (Abcam #ab205720, 1:10,000; Invitrogen #31470, 1:10,000), anti-mouse IgG (H+L) HRP (Promega #W4021, 1:2,500).

### 8. Solubility assays

For sequential solubility analysis, 5 µL of a Percoll-purified schizont stage parasite pellet was lysed in 100 µL of 0.03% (w/v) saponin and incubated on ice for 10 min. The sample was spun down at 16,000 x g for 10 min at 4 °C and washed 2x in ice-cold PBS. The resultant pellet was resuspended in 100 µL of a hypotonic buffer (5 mM Tris-Cl, pH 8.0) and incubated at -20 °C for 30 min (or until completely frozen). The sample was thawed in a 37 °C water bath and centrifuged as before. The supernatant was transferred to a new tube, and the pellet resuspended in 100 µL of 0.1 M Na_2_CO_3_ (pH 11.5) and incubated on ice for 30 min, centrifuged as before, and the supernatant transferred to a new tube. The remaining pellet was resuspended in 100 µL of 1% Triton X-100 and incubated on ice for 30 min, and then centrifuged as before, and the supernatant was transferred to a new tube. The remaining pellet was resuspended in 100 µL of 1x Laemmli buffer and the supernatant in 4x Laemmli buffer (final concentration 1x) and reduced as described before. The fractions were subjected to SDS-PAGE and transferred to a PVDF membrane for Western blot analysis. The method was adapted from Grüring et al., 2012 and Liffner et al., 2020.

### 9. Immunofluorescence assays and co-localisation analysis

#### P. berghei

Protein localisation was assessed using immunofluorescence assays (IFAs) using a previously published method (Jonsdottir et al., 2021) adapted from Tonkin et al., 2004. Whole blood from infected mice with parasitemia below 5% was placed on poly-D-lysine-coated slides and fixed with 4% paraformaldehyde and 0.0075% glutaraldehyde in PBS for 20 minutes at room temperature. Following fixation, cells were permeabilised and quenched using a solution of 0.1% Triton X-100 and 0.1 M glycine in PBS for 15 minutes at room temperature. Blocking was performed with 3% BSA in PBS for 1 hour at room temperature. The samples were then incubated with primary antibodies overnight at 4 °C. After washing, cells were incubated with secondary antibodies for 1 hour at room temperature in the dark. Coverslips were mounted with an anti-fade solution containing DAPI (VECTASHIELD Plus; Vector Laboratories) and sealed with nail polish. Microscopy was conducted using a Leica SP8 inverted confocal microscope equipped with a 63x oil-immersion objective lens. Primary antibodies and concentrations used: rabbit anti-HA (Cell Signal #C29F4, 1:250), rat anti-PbRAP1 (1:500, Proteogenix), and rat anti-PbEXP1 (1:500, Proteogenix). For secondary antibodies, the following were used: Alexa Fluor 647-conjugated goat anti-rabbit IgG (Invitrogen A32733, 1:1,000) and Alexa Fluor 594-conjugated goat anti-rat IgG (Invitrogen A48264, 1:1,000). Image processing, including merging channels, cropping, and adding scale bars, was carried out with ImageJ/Fiji software. The Pearson Correlation Coefficients (Manders et al., 1993) were calculated with a minimum of 3 biological replicates and 60 cells per sample using the JaCoP plug in Bolte & Cordelières, 2006. Graphs were made using GraphPad Prism.

#### P. falciparum

Mature parasites were isolated by Percoll density gradient purification, washed once with PBS, and smeared onto slides and fixed in ice-cold methanol for 5 minutes. Fixed slides were blocked for 1 hour in 3% BSA in PBS followed by overnight incubation with primary antibodies, followed by 3xPBS washes and 1 hour incubation with secondary antibodies in 3% BSA in PBS. Slides were washed 3× with PBS, then Hoechst 33342 (1:500) was added to the sample for 10 minutes, washed 3× with PBS and then mounted with Pro-long Gold mounting solution and cured overnight. Primary antibodies and concentrations used: rat anti-HA (Roche, ROAHAHA, 1:100), mouse anti-PfKAHRP (European Malaria Reagents Repository, Monoclonal antibody 18.2, 1:500), mouse anti-PfRAP1 (European Malaria Reagents Repository, Monoclonal antibody 2.29, 1:1,000), mouse anti-PfEXP2 (European Malaria Reagents Repository, Monoclonal antibody 7.7, 1:1,000). For secondary antibodies, the following were used: Alexa Fluor 488-conjugated goat anti-rat IgG (Invitrogen A11006, 1:500) and Alexa Fluor 594-conjugated goat anti-rat IgG (Invitrogen A11032, 1:500). Visualisation was performed using a Leica LSM880 microscope with a 63x oil-immersion objective and Airyscan processing.

#### Ultrastructure expansion microscopy (U-ExM)

U-ExM was conducted on 4% paraformaldehyde-fixed Percoll-purified schizonts using a protocol adapted from Bertiaux et al., 2021 and Liffner et al., 2023. Briefly, cells were settled onto 12 mm poly-D-lysine coated round coverslips for 20 minutes. Protein anchors were introduced by incubating the cells in a mixture of 1.4% formaldehyde and 2% acrylamide at 37 °C overnight, using a 24-well plate. For gel formation, a solution was prepared by mixing 5 µL of 10% ammonium persulfate and 5 µL of 10% N,N,N’,N’-Tetramethylethylenediamine with a monomer solution containing 23% sodium acrylate, 10% acrylamide, and 0.1% BIS-acrylamide in PBS. A 35 µL drop of this gel mixture was placed onto parafilm, and the coverslips were placed cell-side down onto the gel for polymerisation, and incubated for 1 hour at 37 °C. The gels were then denatured in 1 mL of denaturation buffer (200 mM SDS, 200 mM NaCl, 50 mM Tris in water, pH 9) and incubated at 95 °C for 1 hour and 30 minutes. Gels were expanded by immersing them in 25 mL of double-distilled water (ddH₂O) in 90 mm plates and incubated at room temperature for 30 minutes. The wash was repeated three times, where the gels were left in the last wash to expand overnight at room temperature. The next day, the gels were shrunk by soaking them twice in 1X PBS for 15 minutes. Gels were then placed in 6-well plates and blocked in 2% BSA/PBS for 30 minutes at room temperature or directly probed with 1 mL primary antibody solution diluted in blocking buffer and incubated at RT overnight whilst shaking (125-150 rpm). Gels were then washed three times with 0.1% PBS-Tween, and incubated with secondary antibodies and relevant stains (NHS-Ester, HOECHST, Sytox) in PBS and incubated for 2.5 hours whilst shaking and subsequently washed, as described for primary antibodies. This was followed by three additional washes in 0.1% PBS-Tween. The final expansion was then carried out by soaking the gels three times in 25 mL of ddH₂O for 30 minutes. For gels stained with BODIPY TR ceramide (BODIPY TRc), the stain was diluted in water, and expanded gels were incubated overnight whilst shaking. The next day, the gels were washed three times as before. For imaging, the expanded gels were cut into small pieces and mounted onto 24 mm poly-D-lysine-coated coverslips within an O-ring. Visualisation was performed using a Leica SP8 confocal microscope with a 63x oil-immersion objective. All images were taken as Z-stacks, either using 0.13 or 0.3 µm thick slices, and displayed as maximal intensity projections. Z-projections and adjustments of brightness and contrast were done using Fiji/ImageJ. Primary antibodies used and concentration: rabbit anti-HA (1:250), rat anti-RAP1 (1:250) and EXP1 (1:150). Secondary antibodies used: rabbit anti-405, rabbit anti-488, rabbit anti-647, rat anti-594 all diluted 1:500, Invitrogen. Stains used and concentration: Hoechst 33342 (1:1,000), NHS-488 (1:250), BODIPY TRc (2 µM). Graphs and statistical analysis were done using GraphPad Prism.

### 10. Transmission electron microscopy (TEM)

Samples were fixed in 2.5% Glutaraldehyde (TAAB Laboratories, Aldermaston, England) in 0.1 M PHEM-buffer, followed by post-fixation with 0.8% potassium ferricyanide (K3Fe(CN)6) and 1% osmium tetroxide (OsO4) in 0.1 M PHEM-buffer. Samples were subsequently contrasted with 1% aqueous tannic acid and 1% aqueous uranyl acetate. Dehydration was performed through a graded ethanol series, and samples were embedded in Spurrs resin (TAAB Laboratories, Aldermaston, England) and polymerised overnight at 65 °C. All sample preparation was done using the Pelco Biowave Pro+ processing microwave (Ted Pella, Redding, CA). Ultrathin sections (70nm) were collected on formvar-coated copper grids and analysed using a Talos L120C transmission electron microscope operating at 120kV. Micrographs were acquired with a Ceta 16M CCD camera (FEI, Eindhoven, The Netherlands) using Velox software (version 2.14.2.40 FEI, Eindhoven, The Netherlands). Graphs and statistical analysis were done using GraphPad Prism.

### 11. Co-immunoprecipitation assays

Two mice were infected with WT, RhoSH-HA or MAP1-HA parasite stock via intraperitoneal injection per biological replicate. Once mice had parasitemia of ∼5%, blood was harvested as described above and placed into *ex vivo* culture for 22-24 hours at 37 °C, shaking at 80 rpm. Late schizont stage parasites were purified using 67% Percoll as described above, and the pellet was washed twice in ice-cold PBS containing cOmplete protease inhibitors (Roche) and stored at -80 °C until used. For co-immunoprecipitation assays, pellets were either lysed with triton X-100 or in digitonin. All lysis and immunoprecipitation buffers included protease inhibitors. For the saponin lysis condition, the parasite pellet was lysed in 0.03% saponin in PBS on ice for 10 min and then centrifuged at 16,000 g, and the pellet was lysed in 1% Triton X-100 buffer (150 mM NaCl, 50 mM Tris, pH 7.4). Otherwise, the parasite pellet was directly lysed in 2% digitonin buffer (150 mM NaCl, 50 mM Tris, pH 7.4). For both conditions, samples were freeze-thawed thrice in ethanol/dry ice and spun down at 16,000g for 10 min at 4 °C. Protein concentration of supernatant was measured using Qubit (Thermo Fisher), and 2.5 mg of protein was used per assay. Lysate was made up to 500 µL final volume. Thirty µL of diluted lysate was transferred to a new tube and kept as assay input for western blot. The rest of the lysate was incubated with 60 µL of 50/50 anti-HA agarose bead slurry (pre-washed in immunoprecipitation wash buffer) and let rotate overnight at 4 °C. The next day, beads were washed on ice thrice for 5 min in 1 mL in an immunoprecipitation buffer (Triton X-100 condition) or 0.1% digitonin wash buffer containing protease inhibitors. Beads were then washed 3x in 1 mL PBS, changing tubes after each wash to avoid transfer of detergent. During the last wash, 200 µL (20%) of the solution was transferred to a new tube for western blotting. The rest of the beads were left with just enough PBS to cover the beads and stored at 4 °C for <1 week before elution and analysis using mass spectrometry.

Proteins were then digested by Trypsin/LysC using the iST kit (Preomics) and peptides analysed using nanoLC-ESI-MSMS with the easynLC-1000 liquid chromatography system (Thermo Fisher Scientific), which was coupled with the Q-Exactive HF Hybrid Quadrupole-Orbitrap Mass Spectrometer. All database searches were completed using Mascot (Matrix Science) and the *P. berghei* database and bait protein sequences. Data analysis and validation was completed using the Proteome Discoverer (Thermo Fisher Scientific) using 1% of protein FDR and a minimum of 2 unique peptides per protein.

### 12. Pathology experiments

To evaluate the pathology associated with the KO lines, four C57BL/6 mice were intravenously infected with 10^4 iRBCs. Starting on day 3 post-infection, the mice underwent daily health assessments, including parasitemia measurements via thin Giemsa-stained blood smears, weight monitoring, and clinical scoring. A clinical score was defined based on the presence of the following signs: ruffled fur or piloerection, hunching, and reduced motility. Each category was scored on a scale of 1 (normal) to 3 (severe), with mice being euthanised if their cumulative score exceeded 6, or if they maintained a score of 6 for three consecutive assessments. At the time of euthanasia, spleens were removed and weighed to further assess pathology. To determine the growth rate of the KO lines, parasitemia was calculated from the thin blood smears. For each sample, 1,000 RBCs were counted 100x lens, and parasitemia was expressed as a percentage of iRBCs relative to the total RBCs counted. Graphs and statistical analysis were done using GraphPad Prism.

### 13. *In vivo* imaging (IVIS)

To assess parasite sequestration in organs, IVIS was conducted using a protocol for *P. berghei*, adapted from (Fonager et al., 2012). Briefly, to initiate a relatively synchronous infection, mice received an intravenous injection of 10^6 schizonts purified using Percoll. After 22 hours, mice were anesthetized with isoflurane and administered a subcutaneous injection of 2.4 mg of D-luciferin (Synchem). Imaging was performed 3 minutes after D-luciferin administration using the IVIS Spectrum imager. Imaging sessions were conducted daily at the same time through day four post-infection. On the final imaging day, mice were anesthetized and euthanised to collect spleens for weighing. IVIS images were analysed with Living Image software, ensuring that only images without saturated pixels were used. Regions of interest (ROI) corresponding to the lungs, adipose tissue, and spleen were defined for each mouse during analysis. Graphs and statistical analysis were done using GraphPad Prism.

### 14. Leptin ELISA

Four C57BL/6 mice were subjected to submandibular bleeding and intravenously infected with 10^4 iRBCs. On day 8 post-infection, additional blood samples were collected without anticoagulants. At this time point, the mice were euthanised, and their spleens were harvested. To obtain serum, the blood samples were allowed to clot at room temperature for 30 minutes, then chilled at 4 °C for 5 minutes before centrifugation at 400 × g. The resulting serum was carefully separated and stored at -20 °C until analysis. Serum leptin levels were quantified using a commercial enzyme-linked immunosorbent assay (ELISA) kit (Millipore, EZML-82K; detection range 0.2–30 ng/mL), following the manufacturer’s protocol. For each assay, 5 µL of undiluted serum was used. The experiment was performed in two biological replicates. Graphs and statistical analysis were done using GraphPad Prism.

### 15. Lipidomics

Two mice per biological replicate were infected with mCherry_Luc, Δ*rhosh* or Δ*map1* parasite stock via intraperitoneal injection. Once parasitemia reached around 5%, blood was collected via cardiac puncture, and the blood from both mice was pooled together and incubated *ex vivo* for seven hours. Trophozoite stage parasites were harvested using 67% Percoll density gradient as described above and washed twice in RPMI before being placed in cryotubes and snap frozen on dry ice. Lipid extraction and analysis were performed following a biphasic extraction protocol optimised for LC-MS/MS. LC-MS grade water, acetonitrile (ACN), methanol (MeOH) and isopropanol (IPA) were obtained from Th. Geyer (Germany). High-purity methyl tert-butyl ether (MTBE), ammonium formate, formic acid, ammonium acetate, and acetic acid were purchased from Merck (Germany). Stable isotope-labelled internal standards (EquiSPLASH; Avanti Polar Lipids, AL, USA) were used at final concentrations of 0.5% (v/v).

For metabolite extraction, 1 mL of MeOH containing internal standards was added to cell pellets and homogenised on dry ice using a bead beater (FastPrep-24; MP Biomedicals, CA, USA) at 6.0 m/s (3 × 30 s, 5 min pause) with 1.0 mm zirconia/glass beads (Biospec Products, OK, USA). After centrifugation for 10 min at 15,000 × g and 4 °C , supernatants were transferred to 5 mL tubes, and 2.5 mL of MTBE were added. The monophasic mixture was vortexed for 60 s and incubated at −20 °C for 20 min, followed by the addition of 625 µL water, vortexing, and a second incubation under the same conditions. Phase separation was achieved by centrifugation for 10 min at 15,000 × g and 4 °C. For lipidomics analysis, 2 mL of the upper organic phase were collected, dried under a nitrogen stream, and reconstituted in 100 µL of isopropanol:methanol (50:50, v/v).

Chromatographic separation and detection were carried out on a Vanquish UHPLC system coupled to an Orbitrap Exploris 240 high-resolution mass spectrometer (Thermo Scientific, MA, USA), operated in both positive and negative ESI modes. For untargeted lipidomics, analytes were separated on an ACQUITY Premier CSH C18 column (Waters, 2.1 × 100 mm, 1.7 µm) at 0.3 mL/min, with mobile phases consisting of water:ACN (40:60, v/v; phase A) and IPA:ACN (9:1, v/v; phase B), modified with 10 mM ammonium acetate + 0.1% acetic acid (negative mode) or 10 mM ammonium formate + 0.1% formic acid (positive mode). The 23-min gradient was as follows (min/%B): 0/15, 2.5/30, 3.2/48, 15/82, 17.5/99, 19.5/99, 20/15, 23/15. Column temperature was maintained at 65 °C, the autosampler at 4 °C, and the injection volume was 5 µL. Full-scan data were acquired at 120,000 resolving power (200–1700 m/z, 100 ms scan time), and data-dependent MS/MS spectra were obtained at 15,000 resolving power using stepped collision energies of 25/35/50%. Ion source parameters were set to the following values: spray voltage: 3250 V / -3000 V, sheath gas: 45 psi, auxiliary gas: 15 psi, sweep gas: 0 psi, ion transfer tube temperature: 300°C, vaporizer temperature: 275°C.

All samples were measured in randomised order. Quality control (QC) samples were prepared by pooling equal aliquots of all processed samples. Multiple QC injections were performed at the start of the sequence for system equilibration, and additional QCs were analysed every fifth sample to monitor instrument performance. A processed blank was included for background subtraction. Data were processed with MS-DIAL 4.9.221218 (Tsugawa et al., 2015), and peak intensities were normalised to the total ion count of detected features (Drotleff & Lämmerhofer, 2019). Level-1 metabolite identification was achieved using the MS-DIAL LipidBlast V68 library, based on accurate mass, isotope pattern, MS/MS fragmentation, and retention time matching, with a minimum score threshold of 80%.

For analysis of lipidomics data, for each sample, the relative abundance of every lipid species was calculated by normalising the intensity of each lipid species to the total lipid intensity within that sample (i.e., the sum of all lipid species per sample equals 1). This normalisation allows direct comparison of lipid abundances between the mCherry-Luc control, Δ*rhosh*, and Δ*map1*, with four biological replicates analysed per parasite line. To assess global changes at the lipid headgroup level and capture any lipid classes affected by gene knockout, relative abundances of individual lipid species belonging to the same headgroup were summed, providing the overall relative contribution of each headgroup within a sample. Statistical analysis was performed using one-way ANOVA (implemented via the scipy.stats.f_oneway function) to identify significant differences between groups. For each lipid headgroup, p-values and log₂ fold changes were calculated. Statistical significance was set at *p* < 0.05 with a log₂ fold change cut off of ≥ 0.58 or ≤ −0.59. Visualisation of results was conducted in Python using the plotnine package. Volcano plots were generated to illustrate the relationship between fold changes and statistical significance, while boxplots were used to display the distribution of summed relative intensities across conditions.

### 16. CD36 binding assay

Assay was adapted from Kundu et al., 2023. Twenty µL streptavidin coated fluorescent Nile red particles 0.4-0.6 µm (Spherotech SVFP-0556-5) were blocked in 500 µL 1%BSA in PBS for 30 minutes at room temperature whilst rotating, centrifuged at 9,400 × g for 5 minutes and resuspended in 20 µL 1%BSA in PBS. The blocked beads were sonicated for 10 minutes (30s on, 30s off 4 °C 10 cycles using a Diagenode Bioruptor Pico) and conjugated to biotinylated CD36 (Acrobiosystems CD6-H82E9) for 45 minutes on ice with a ratio of 1 µL of beads to 1 µg biotinylated CD36, multiplied by the number of planned assay wells. The coated beads were washed twice with 500 µL 1% BSA in PBS. Centrifugation steps were carried out at 9,400 × g for 5 minutes at 4 °C. After the final wash, the beads were resuspended in 1%BSA in PBS at 1 µL per planned assay well.

Late-stage parasites, which had been sorbitol synchronised and cultured for two cycles ± 2.5 mM GlcN, were isolated on a 63% Percoll column, washed and resuspended in PBS plus sybr green 1:5,000 at 37 °C for 30 minutes, washed and resuspended in 1 mL complete media (± GlcN). Cells were counted using a C-chip disposable haemocytometer and 5 × 10^5 cells were added to triplicate assay wells with 1 µL of conjugated beads in a total volume of 200 µL complete media (± GlcN) per well and incubated for 30 minutes at room temperature. Three washes were carried out with all centrifugation steps at 190 × g for 10 minutes at room temperature to remove unbound beads.

All wells were analysed by flow cytometry, and gates were applied to separate the population of Nile red-positive and Sybr green-positive cells. Binding efficiency was expressed as a percentage of control wells, and an unpaired t-test was carried out using Graphpad Prism.

For the PfMAP1-3xHAglmS transfectant, three biological replicates were carried out, each with three technical replicates. For 3D7 WT parasites, only two biological replicates were carried out with a total of four assay wells between them due to limiting amounts of CD36 protein.

## Figures

All figures and schematics were made and arranged using Adobe Illustrator.

## Acknowledgments

We gratefully acknowledge Umeå Centre for Electron Microscopy (UCEM) and the Biochemical Imaging Center at Umeå University for their support with microscopy. Additionally, we thank the University of Geneva Proteomics Facility for their assistance with proteomic analyses and data interpretation. We acknowledge the support of the EMBL Metabolomics Core Facility (MCF) in the acquisition and analysis of liquid chromatography-mass spectrometry data. We thank Paul R. Gilson and Dawson B. Ling (Burnet Institute) for providing the modified pSLI vector. We thank Prasun Kundu (Cambridge Institute for Medical Research) for giving advice on the *Plasmodium falciparum* iRBC CD36 binding assay.

## Author Contributions

M.S.P., T.K.J. and E.S.C.B. conceptualised the study and wrote the original manuscript. M.S.P., T.K.J., A.K, S.A.C, S.H. and M.R.D planned and conducted experiments. M.S.P., T.K.J., A.K, S.A.C, S.H., C.Y.B. and V.P. analysed data. A.B. and R.P.A.B performed the AlphaFold analysis and made relevant graphs. T.K.J. put together the final figures. E.S.C.B., R.F.W. R.P.A.B and J.C.R. supervised. All authors participated in the review and editing of the final manuscript.

