## Supplementary Information for "Characterisation of Two *Plasmodium* Virulence Factors Important for Lipid Metabolism and Disease Progression *In Vivo*"

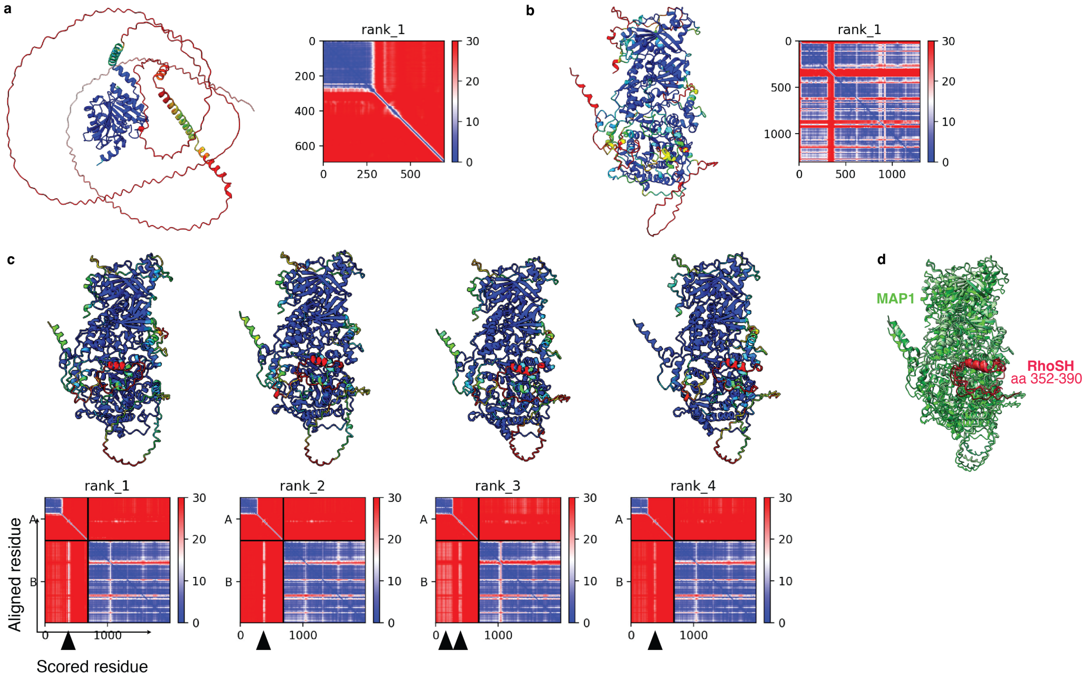


**Figure S1. Alphafold2 predictions of MAP1 and the RhoSH-MAP1 complex.**
AlphaFold2 models of **a** RhoSH (rank 1) and **b** MAP1 (rank 1), coloured according to the pLDDT score (predicted local distance difference test), with the corresponding predicted aligned error (PAE) plots. **c** AlphaFold2 models rank 1-4 of the RhoSH-MAP1 complex, generated using the RhoSH sequence and the predicted MAP1 (rank1) model as input. The models are coloured according to pLDDT score: corresponding PAE plots are shown below. Predicted interaction sites are indicated by black triangles. PAE plot: Chain A corresponds to RhoSH and chain B to MAP1. **d** Superimposition of all AlphaFold2 models highlighting the similar predictions for the RhoSH-MAP1 complex. The models of MAP1 are shown in shades of green and the hydrophobic peptide side chain of RhoSH in shades of red.

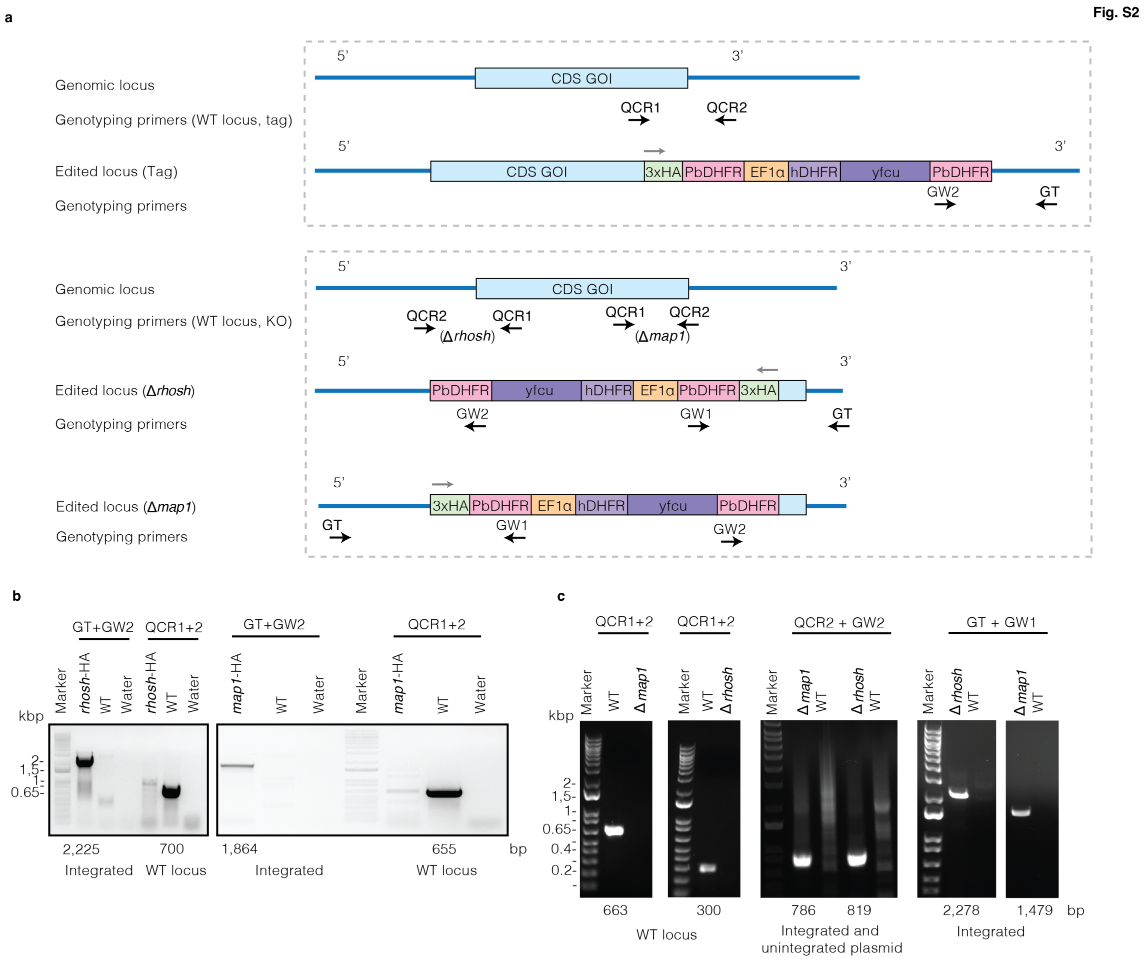


**Figure S2: Genotyping PCRs for *rhosh* and *map1* tagging and KO *P. berghei* transgenic parasite lines.**
**a** Schematic overview of where genotyping primers are located for *rhosh* and *map1* tagged/knockout parasites using the *Plasmo*GEM system (Gomes et al., 2015) with primer sequences obtained from https://plasmogem.serve.scilifelab.se/pgem/home. GT = genotyping (upstream or downstream of homology region), QCR1 = Quality Control R1 (within gene of interest (GOI) coding DNA sequence (CDS)), QCR2 = Quality Control R2 and GW = Gateway (vector backbone) **b** Genotyping PCRs for *rhosh* and *map1* tagged with 3xHA. **c** Genotyping PCRs for Δ*rhosh* and Δ*map1*.

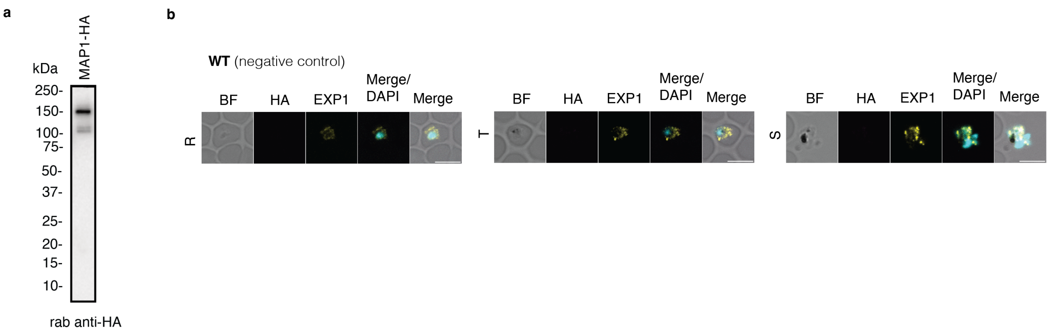


**Figure S3: Western blot for MAP1-HA and immunofluorescence assays for WT control parasites.**
**a** Western blot for MAP1-HA probed with rabbit anti-HA antibody, where signal is observed at the expected size with additional two processed forms of MAP1-HA around 100 kDa. **b** Immunofluorescence assays of *P. berghei* WT parasites, where rabbit anti-HA showed no signal and rabbit anti-EXP1 observed at the PVM as expected. Scale bars = 5 $\mu$m.

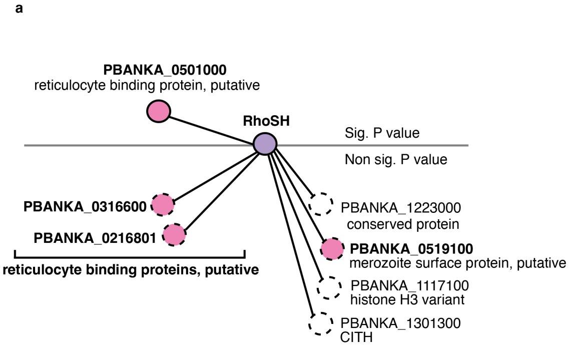


**Figure S4: Co-immunoprecipitation assays of RhoSH-HA and MAP1-HA saponin lysed and Triton X-100 treated parasite pellets.**

Co-immunoprecipitation assays were completed on **a** RhoSH-HA and **b** MAP1-HA, where schizont pellets were lysed in saponin and subsequently lysed in Triton X-100. The horizontal line marks what was considered significant interactions compared with WT parasite as a control. Target proteins are indicated in purple and protein of potential interest in pink. Proposed interaction network only shows proteins with significant interaction or enriched but not significant (indicated by the horizontal line) compared to the WT control from three biological replicates.

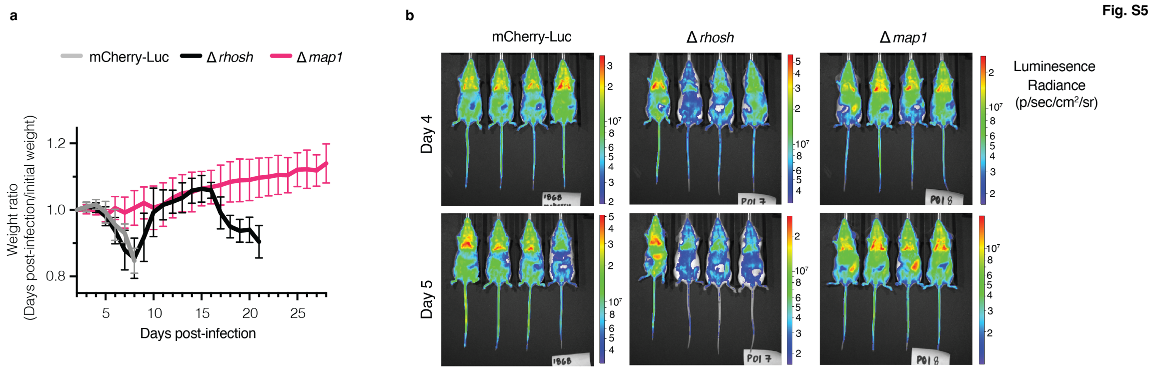


**Figure S5: Additional data for pathology and IVIS experiments presented in Fig. 4.**
**a** All mice used for pathology experiments in **Fig. 4a-c** were weighed at the start of the experiment and until the end point to ensure mice were within acceptable range (<20% weight loss, with mice exceeding 20% weight loss euthanised). Error bars = standard deviation for three individual experiments, where each experiment included four mice per condition. **b** IVIS images accompanying **Fig. 4d** graphs for day 4 and 5.

Fig. S6

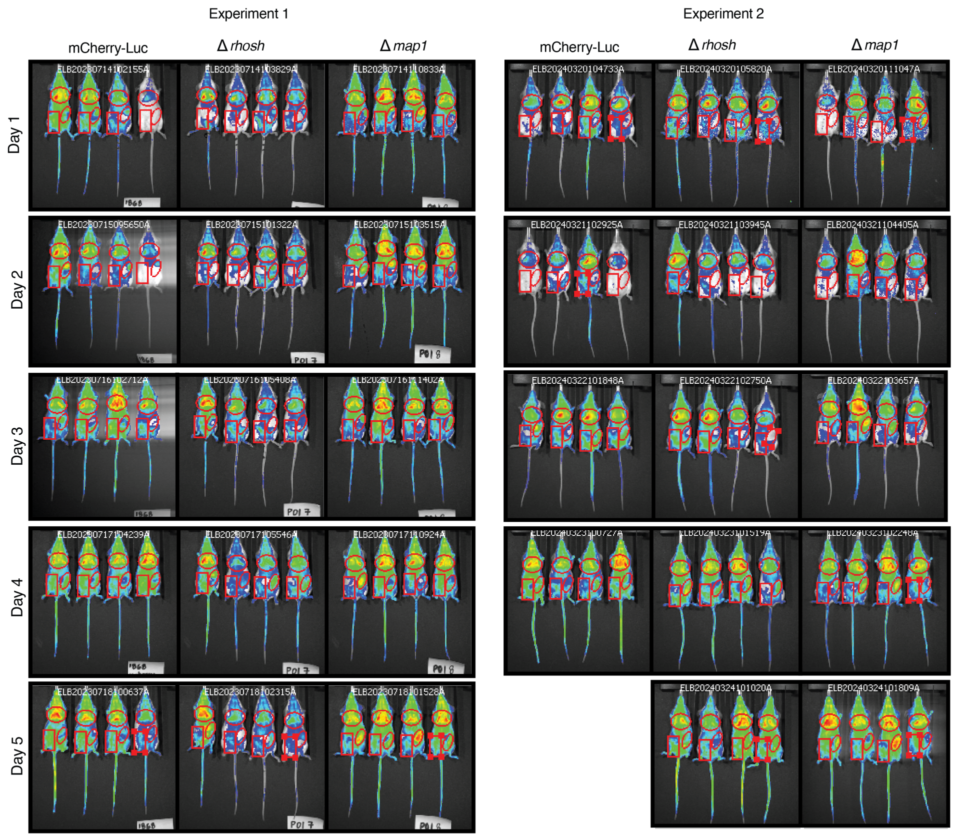


**Figure S6: IVIS experiments in Fig. 4d with regions of interest labelled.**

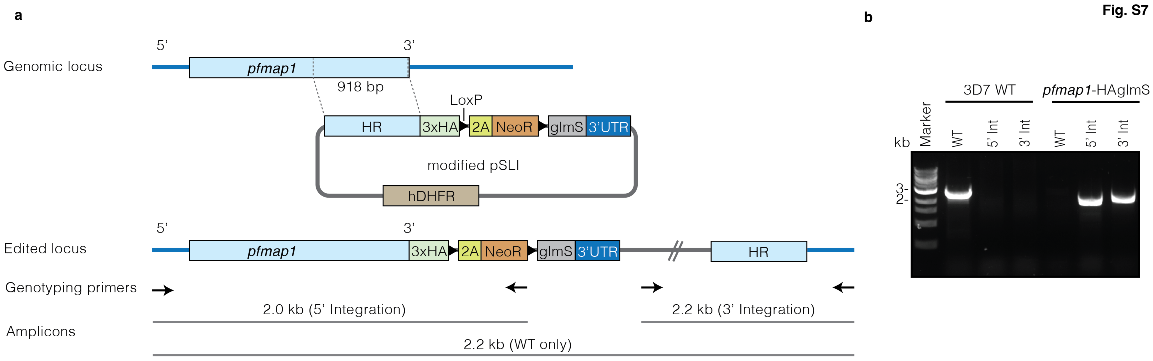


**Figure S7: The generation and confirmation of the PfMAP1-HAglmS parasite line.**
**a** To generate the *pfmap1*-HAglmS line, 918 bp homology region (HR) was chosen just before the gene stop codon and ligated into the modified pSLI vector in frame with 3xHA. Genotyping primers and expected amplicon sizes post integration are indicated in relevant places. **b** Genotyping PCRs for *pfmap1*-HAglmS parasites, where the genotyping primers and expected amplicon sizes post integration are indicated in relevant places on the schematic on the left side.

**Table S1. Primers used for generation and genotyping of *P. berghei* and *P. falciparum* mutant lines.**

| Primer name | Sequence (5' -> 3') | Note |
|---|---|---|
| PB_1001500_QCR1_tag | TGCGTCGAAGAGTGATGCCCT | PlasmoGEM primer for tagging |
| PB_1001500_QCR2_tag | GCGTGTGCTGTCATTACCACT | PlasmoGEM primer for tagging |
| PB_1001500_GT | TGCTGCCAGCTACTCGCTTACCT | PlasmoGEM primer for tagging and KO |
| PB_1001500_QCR1_KO | TTGCATAACTCGGTGGATGTGGGC | PlasmoGEM primer for KO |
| PB_1001500_QCR2_KO | AAACGCGCAGAATGGCTACA | PlasmoGEM primer for KO |
| PB_1425900_QCR1_tag | GCATCAGCCAAACCCACACTTGG | PlasmoGEM primer for tagging |
| PB_1425900_QCR2_tag | ACGTGAACGAATCGTGTAATTGT | PlasmoGEM primer for tagging |
| PB_1425900_GT_tag | TGCGAACAATCCAGAAAGCACCG | PlasmoGEM primer for tagging |
| PB_1425900_QCR1_ko | TTTGTGCACGCAGATGCATT | PlasmoGEM primer for KO |
| PB_1425900_QCR2_ko | CCAAGTGTGGGTTTGGCTGATGC | PlasmoGEM primer for KO |
| PB_1425900_GT_ko | TTTGTGCACGCAGATGCATT | PlasmoGEM primer for KO |
| GW1 | CATACTAGCCATTTTATGTG | PlasmoGEM generic primer |
| GW2 | CTTTGGTGACAGATACTAC | PlasmoGEM generic primer |
| pSLI_Fw_081 | GCTAACGTAACAGACTTAGGAGGAGATC TGCATGATGATGATATTAGTCTTGGTT | Amplification of homology arm with Gibson overhang for vector construction |
| pSLI_Rv_081 | CCGGGACGTCGTACGGGTAAGAGGCTG CAGCTTCGTCTACCTTAAATAAATAAAAG AAG | Amplification of homology arm with Gibson overhang for vector construction |
| PF0811600_5'int_Fw2 | CGGATAATGTTAATATGAATATCGTAGA C | Flanking primer forward for genotyping |
| PF0811600_3'int_Rv1 | GTACCAAATGTTTCAAATAATACAGGTAC | Flanking primer reverse for genotyping |
| NeoR_Nterm_5INT_R | CCCAAGCAGCTGGTGATCC | Universal SLI genotyping primer |
| pSLI_3INT_F | TTGGCCGATTCATTAATGCAGC | Universal SLI genotyping primer |
